# Common organization of collective dynamics across divergent self-organized neural architectures

**DOI:** 10.64898/2026.11.18.767733

**Authors:** Zehui Zhao, Liang Yang, Ilya Nemenman, Andre Levchenko

## Abstract

Brain organoids and other model neuronal networks build synaptic circuits through self-organization, and different protocols yield markedly different architectures, letting us test how collective dynamics depend on large-scale organization. We compared organoids of layered cortical-like structure against networks of cell-body clusters joined by axonal cables, quantifying the reproducible patterns of population activity in each. Bursts occurred in both architectures and comprised several collective patterns in each, so no single recurring template describes them. Recurrent activation sequences capture only part of this organization, sequence participation is graded, and neurons that rarely join a sequence still contribute to the shared dynamics. The patterns also pick out overlapping neuronal groups with propagation-like spatial order. Although the patterns differed between tissue models, their organization was conserved, and remained unchanged under pharma-cological increases in network activity. Recurrent sequences therefore represent one manifestation of a broader collective organization that emerges across distinct self-organized neural architectures.

## I. INTRODUCTION

How the dynamics of neural populations depend on the architecture of the circuits generating them remains a central problem in neuroscience. Across many systems, activity organizes into reproducible population patterns, among them propagating waves [1, 2], low-dimensional trajectories [3, 4], and activation in a stereotyped temporal order [5, 6]. Whether such patterns require particular anatomical arrangements, or arise wherever neural tissue intrinsically self-organizes, is unclear.

Model neuronal networks, including brain organoids (BOs) derived from human induced pluripotent stem cells (hiPSCs), allow this dependence to be tested directly. These model systems generate active synaptic circuits through developmental self-organization, which reproduces the diverse neuronal cell types, regional patterning, and tissue architecture of developing human neural tissue, albeit in a simplified manner [7–16]. Detailed connectivity among individual neurons is left unspecified by the experimental conditions and emerges through developmental and activity-dependent processes [17], so that microscopic wiring can vary even where gross neural organization is apparently similar.

BOs (Figure 1A) recapitulate the key characteristics of early brain development, including progenitor organization and neuronal stratification, although the full layered architecture of the cortex is not reproduced [8, 9, 15, 18]. Alongside them, other protocols guide neurogenesis and model tissue organization into localized neuronal clusters joined by long axonal bundles, which we here refer to as *cable-cluster* networks (CCNs, Figure 1A). CCNs display an architecture more similar to ganglia or brain nuclei connected by tracts, i.e., cables of fasciculated nerve fibers found in both the central and peripheral nervous systems [19, 20]. Engineered neuronal tissues such as CCNs often require a scaffold, whereas BOs generate their own three-dimensional organization through intrinsic developmental programs. The two architectures differ most clearly at the 400–500 *µ*m mesoscale over which population dynamics are commonly measured, where a BO presents a continuous, highly interconnected and integrated population of neurons, much like the cerebral cortex in vivo, and a CCN is organized into several spatially separated neuronal aggregates. These ganglion-like clusters of cell bodies lack the layered patterning of cortical tissue, as their relatively homogeneous distribution of neuronal markers shows, while their constituent cells remain strongly coupled [19, 20]. Within a cluster, cellular composition shares much with both in vivo developing neural tissue and BOs [19, 20], so the two cytoarchitectures differ mainly at intermediate and large scales of cellular organization while sharing cell type makeup. Comparing them therefore isolates what large-scale cell arrangement contributes to the dynamics.

**FIG. 1.**
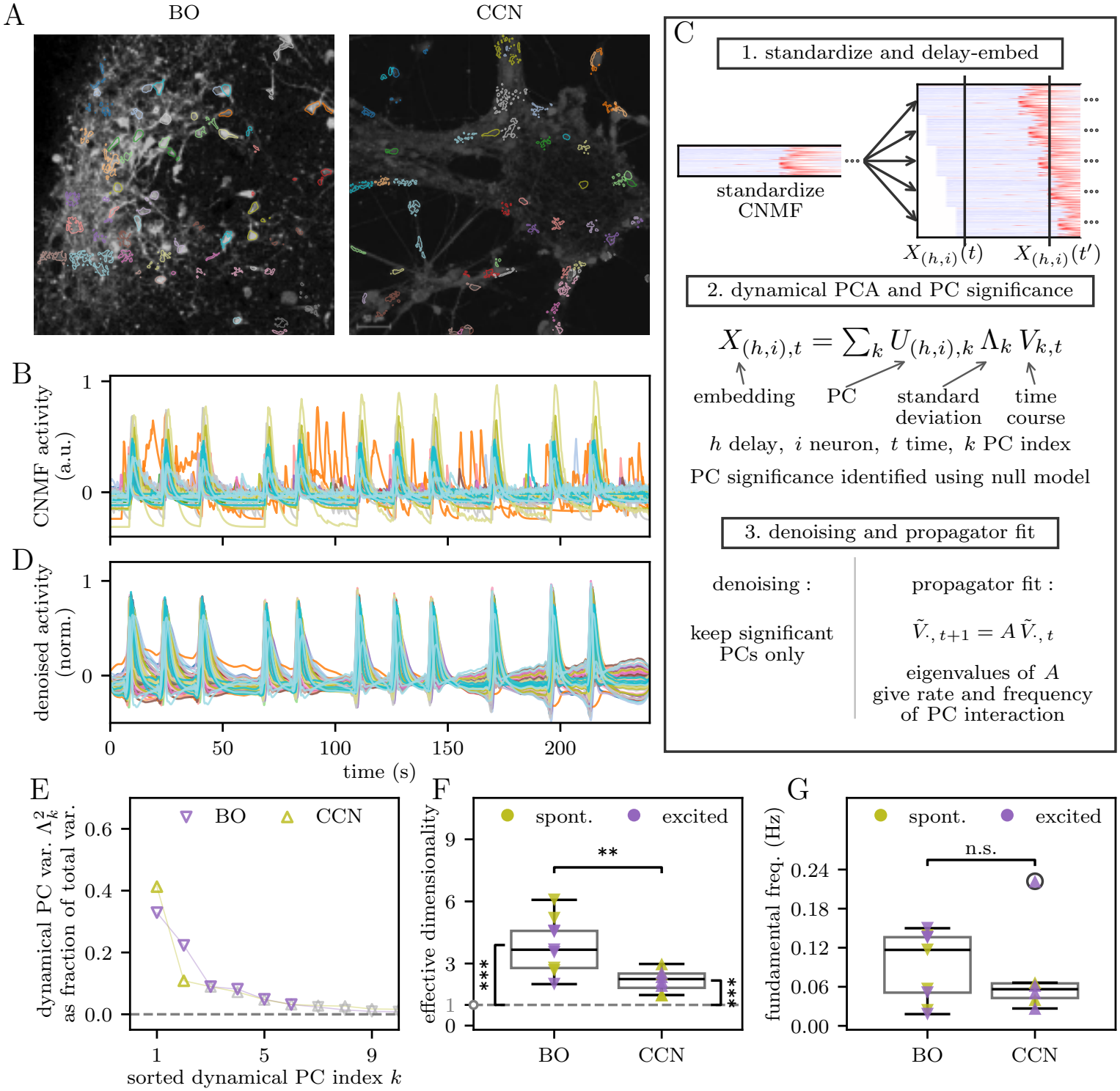
Collective Patterns and Their Rate Across Cytoarchitectures. A: Sample fluorescence snapshots with CNMF-identified neuron contours for calcium activity in a BO slice following bicuculline treatment (left) and in a spontaneously active CCN (right). B: CNMF activity for the BO slice shown in A, plotted over time with one color per neuron and scaled by its maximum. C: Analysis pipeline: (1) standardize and delay-embed CNMF activity into a stacked matrix; (2) perform principal component analysis (PCA) and compare singular values against a matched surrogate null; (3) fit a one-step propagator on the trajectories of the significant dynamical principal components (PCs), whose eigenvalues give each dynamical mode a frequency. D: The denoised activity in B reconstructed from its significant dynamical PCs, on the same time axis and scaled by its own maximum because the denoised activity is in standardized units. E: Fraction of total variance carried by low-index dynamical PCs, against sorted dynamical PC index, for the BO recording of A, B and D and the CCN recording of Markers follow the figure conventions; gray markers denote dynamical PCs within the range of that recording’s surrogate spectrum and colored markers denote the significant dynamical PCs outside that range. Significant dynamical PCs account for the majority of embedded variance in both. F: Effective dimensionality for BO and CCN calcium activity, estimating the number of significant population patterns after downweighting low-variance dynamical PCs. Both cytoarchitectures support multidimensional dynamics, with BOs showing a larger repertoire. The dashed horizontal line marks the reference value 1; the vertical bracket beside each group compares that group with the reference. G: Fundamental frequency, the lowest oscillation rate resolved by the fitted propagator, for BOs and CCNs. Recordings resolving no oscillation above the two-cycle floor are omitted. The rate of collective activity is a shared feature across cytoarchitectures. Statistics: F,G, unpaired Welch t-tests for BO vs. CCN comparisons, shown as horizontal brackets; F, Holm-corrected one-sample one-sided t-tests against 1, shown as vertical brackets. **Figure conventions**. For summary plots, points denote individual movies unless otherwise stated. Olive and purple denote spontaneous baseline and pharmacologically excited conditions, respectively; downward and upward triangles denote BOs and CCNs, respectively. Asterisks: * *p ≤* 0.05, ** *p ≤* 0.01, *** *p ≤* 0.001, **** *p ≤* 0.0001.

**FIG. 2.**
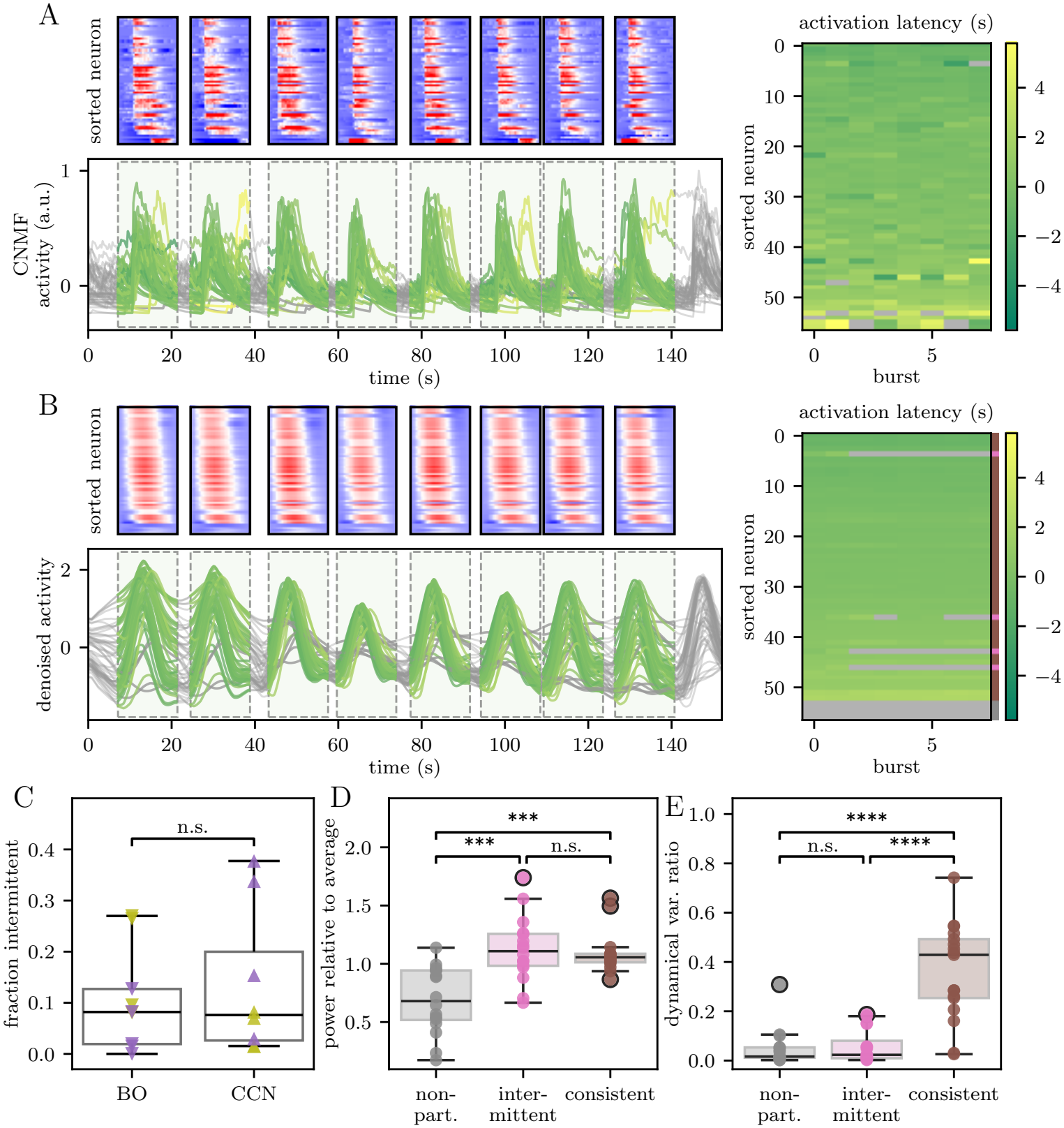
Neurons Join the Sequence Intermittently While Remaining Part of Collective Dynamics. A: CNMF-based activation-sequence analysis before denoising for an example spontaneous CCN recording: top, burst-aligned heatmaps sorted by mean activation time; bottom, normalized traces; right, burst-by-neuron latency matrix. Heatmaps show endpoint-detrended CNMF activity on one common color scale after normalizing the recording to its maximum: zero and negative detrended values are blue, and increasing positive activity progresses toward red. They carry no residual-noise participation threshold. B: Activation-sequence analysis after denoising for the same recording: top, burst-aligned heatmaps sorted by mean activation time; bottom, denoised traces; right, burst-by-neuron latency matrix. In A and B, each heatmap is positioned directly above its corresponding shaded burst window, with its left and right edges marking the start and end of that window. Heatmap color gives detrended denoised activity divided by the neuron’s residual-activity standard deviation on one scale across all bursts; blue lies below the participation-amplitude threshold, white marks the threshold at one residual standard deviation, and red lies above it. Each burst column in the right panels follows the same neuron order, and latency is each neuron’s activation time relative to the other neurons in that burst. Green traces: sequence-participating activation; gray traces: non-participating; burst windows are shaded; the latency matrix is colored by activation latency. The narrow bar to the right of the denoised latency matrix labels each neuron by sequence participation, colored as in D and E: gray, non-participating; pink, intermittent; brown, consistent. Denoising keeps only the latency assignments supported by significant dynamical components, and membership in the sequence varies from burst to burst. C: Fraction of neurons classified as intermittent in each BO and CCN recording. Intermittent neurons occur in nearly every recording and comprise similar fractions of the two cytoarchitectures. D: Variance-weighted normalized squared loading across recordings, by participation type. Non-participating neurons have lower normalized squared loading than intermittent and consistent types, but all three are significantly above zero. E: Dynamical variance ratio across recordings, by participation type. Consistent neurons express more burst-timescale variation in collective activity on average than non-participating or intermittent types, while varying widely across recordings. Statistics: C, unpaired Welch t-test comparing BOs and CCNs; D,E, one-way ANOVA across participation type, factorial ANOVA for participation type by cytoarchitecture interactions, Holm-corrected unpaired Welch t-tests between types, and Holm-corrected one-sample one-sided t-tests against zero; E, Brown–Forsythe test comparing spread (*W* = 17.7, *p* = 1.2*×* 10^−4^). In plots involving sequence participation, gray, pink, and brown denote non-participating, intermittent, and consistent neurons.

**FIG. 3.**
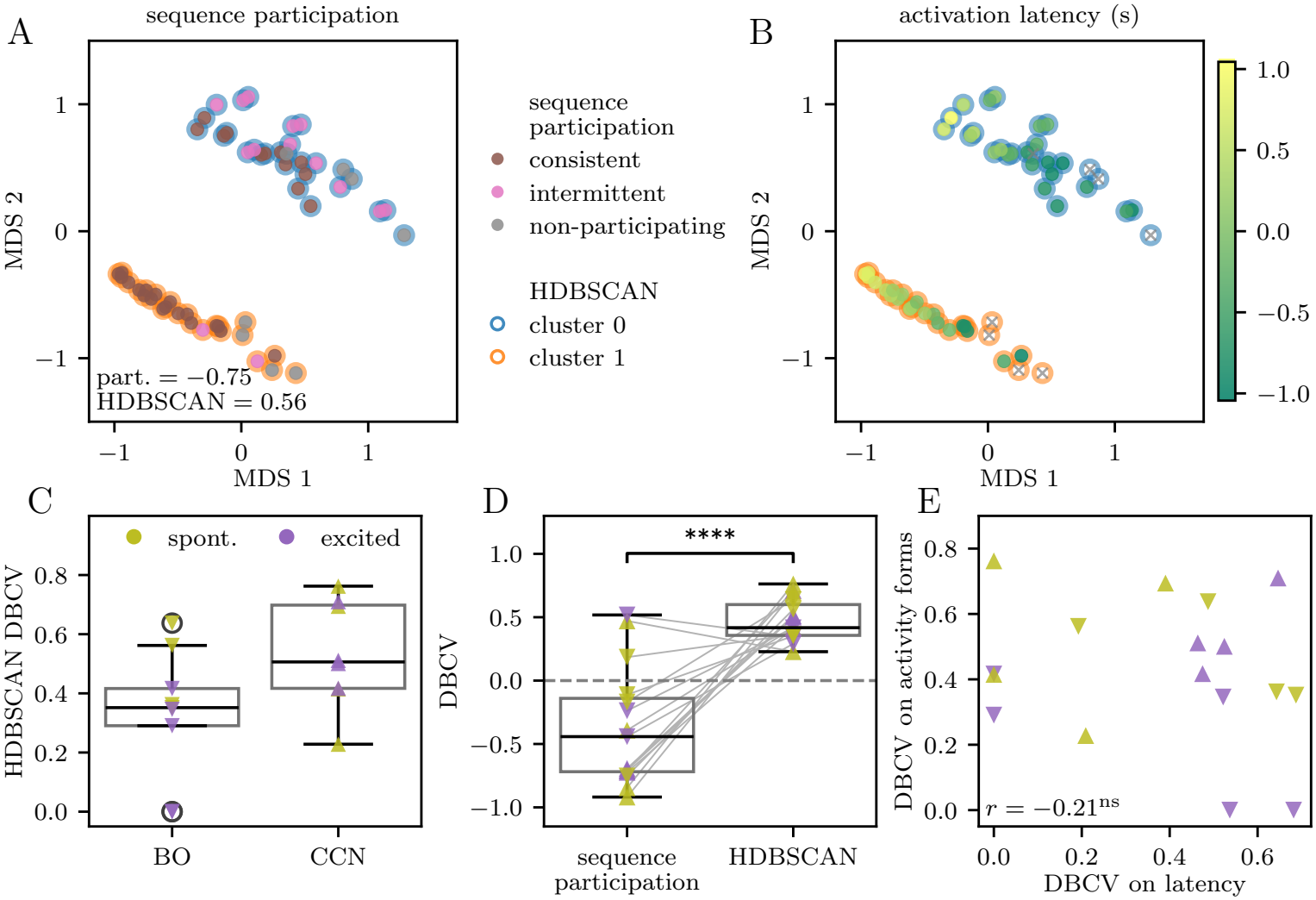
Neurons Come in a Few Activity Types, Unrelated to the Sequence. Throughout, clustering is quantified by the DBCV score, positive when neurons are more similar within groups than between them and negative otherwise. A: Multidimensional scaling (MDS) embedding of activity-form distances for an example spontaneous BO recording; fill color: participation type, halo color: activity type (HDBSCAN); number: DBCV of this recording. Nearby points have similar activity forms, normalized profiles within the significant dynamical principal components (PCs). Each cluster mixes all three participation types. B: Same embedding, colored by activation latency; gray x markers: non-participating; halos mark activity types. Latency varies within clusters without separating them. C: DBCV for BOs and CCNs. Cluster separation is similar in both cytoarchitectures. D: DBCV of the same activity-form distances under participation labels or activity type labels; gray lines connect scores from the same recording; dashed line marks zero. Cluster labels score positive and participation labels negative, so participation types overlap in activity form. E: DBCV of activity-form vs. latency-profile clustering. The two are uncorrelated, so activity types do not follow activation timing. Statistics: C, unpaired Welch t-test; D, paired one-sided t-test, with Holm-corrected one-sample tests against zero; E, Pearson correlation.

**FIG. 4.**
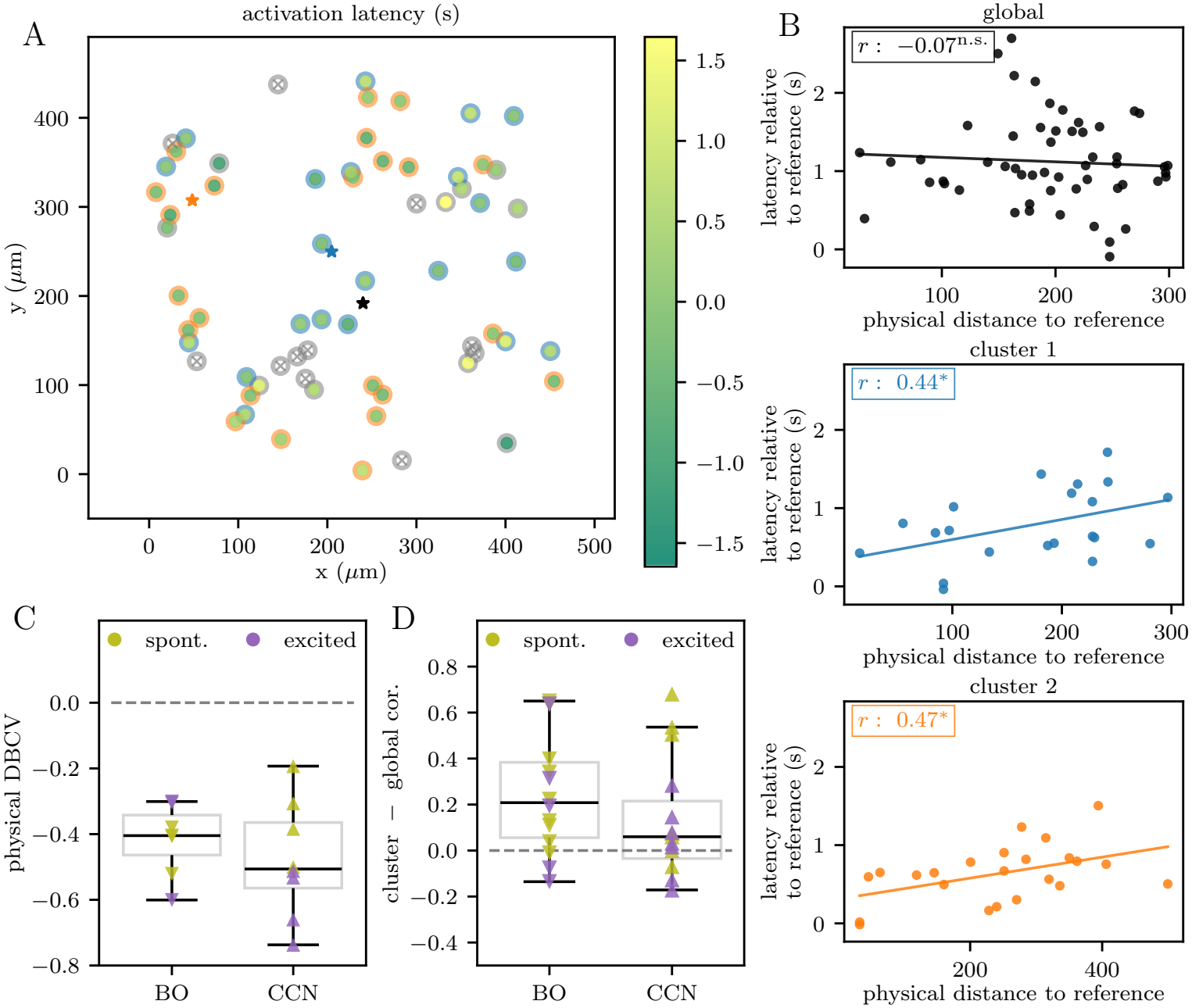
Activity Types Overlap in Space, and Activity Spreads Within Each of Them. A: Physical positions of neurons in the spontaneous CCN recording with the largest within-type gain in D, colored by latency; gray x markers: non-participating; colored halos: activity types, gray halos: unassigned neurons. Stars: early-latency centroids for the full population and each type. The types are mixed in space. B: Latency vs. physical distance, both relative to the early-latency centroid, for the full recruited population and within each type. Lines: linear fits; numbers: Pearson correlation and significance. The correlation is stronger within types. C: DBCV of activity types on physical distances, BOs and CCNs. Negative values: the types are not segregated in space. D: Within-type minus whole-population distance–latency correlation, one point per activity type, BOs and CCNs. Across recordings the correlation is higher within types than for the whole population; the increase is significant in BOs, not in CCNs alone, and does not differ detectably between the two. Statistics: B, Pearson correlations; C, D, Welch t-tests between cytoarchitectures and Holm-corrected one-sample tests against zero within each (two-sided in C, one-sided in D), plus a pooled one-sided test in D.

Brain organoids and related model networks are increasingly used to study development, disease, and the action of drugs, and these are often read from their effect on the neuronal population activity. That effect, however, is usually measured by a single summary statistic, such as firing rate, burst frequency, or synchrony. In the brain, population activity, spontaneous activity included, is far richer than any such statistic: it unfolds along many dimensions at once, and its structure, more than its overall level, is what distinguishes one state of a circuit from another [4, 21]. Organoid activity is rarely described at this level, and a description of comparable resolution is needed for an organoid to report what a change in development, a disease, or a drug does to its circuits.

Indeed, it is known that spontaneous bursts occur in BOs and other model neuronal networks as reported by us and others [14, 22, 23]. It was argued that these bursts contain recurrent activation sequences, in which neurons become active in a reproducible relative order across repeated events, and that the sequences are carried by a subset of cells defined as a “backbone” [6, 23]. Neither sensory experience nor designed connectivity is available or needed to produce them, placing their origin in neuronal self-organization and making these preparations a test of what developmental processes alone can build. However, several points remain unresolved. First, a reproducible order establishes the existence of temporal structure, but it leaves open whether that structure amounts to a single recurring pattern or reflects a richer underlying repertoire of collective dynamics. Second, the geometric and anatomical organization (topography) of these distinct collective patterns in physical space remains unknown. Resolving this question is particularly relevant to understanding how recurrent bursting activity can organize the maturation of connectivity in developing neural systems. Third, backbone participation could mark a discrete division between sequence-generating neurons and the rest, or that division could be arbitrary, with participation instead grading continu-ously into a broader collective organization of neuronal activity. Finally, if sequential activity depends strongly on cytoarchitecture and on the relative composition of neuronal types, qualitatively different tissue organizations would be expected to produce qualitatively different dynamics. Conversely, if sequence generation reflects more general principles of neural self-organization, similar dynamics may emerge across anatomically distinct tissues. Distinguishing among these possibilities is necessary to determine how strongly collective neural dynamics are constrained by the individual neuronal components and the network architecture.

Here we address these questions by comparing population activity across two distinct self-organized neural architectures: cortical-like BOs patterned by orthogonal morphogen gradients in the DuoMAPS device [14], and CCNs generated by classical dual-SMAD inhibition with a WNT inhibitor, as previously described [16]. To accomplish this, we analyzed time-delay-embedded population activity and retained only the activity patterns significant compared to matched surrogate null model. Despite their anatomical differences, both cytoarchitectures exhibited similarly organized collective dynamics, with burst activity resolving into several recurrent patterns rather than a single dominant one. Recurrent sequences contribute substantially to these dynamics without fully defining them, and participation in the sequences is graded, depending on which collective patterns a neuron contributes to and how strongly those patterns are expressed. The collective patterns reveal overlapping functional subnetworks with propagation-like spatial organization. These features persist despite pharmacological elevations of network activity, suggesting that they reflect intrinsic properties of the underlying circuitry. Distinct self-organized architectures therefore converge onto a common organization of collective dynamics, of which recurrent activation sequences are but one manifestation.

## II. RESULTS

### A. Bursts Contain Several Collective Patterns in Both Cytoarchitectures

BOs and CCNs develop without sensory experience or designed connectivity (Figure 1A), yet both produce coordinated spontaneous bursts, making this activity selforganized [23]. Prior work described these bursts as noisy repetitions of a single dominant pattern, a fixed template [23]. But is one pattern all a burst contains? And does the answer depend on the large-scale cytoarchitecture, or on the state of the network, such as its typical activity rate?

To answer these questions, we examined the calcium readouts as a proxy for electrical cell activity, by segmentation of the movies to extract the activity of individual neurons. We then analyzed the temporal covariation of this activity, which was not necessarily simultaneous, since neurons in a burst can activate at different times, keeping only the components of that covariation that exceeded what finite sampling and single-neuron autocorrelation produced on their own. Finally, we sought to understand how these components arrange themselves into dynamical modes that evolve in time with definite rates and frequencies. Each of these properties can then be compared across cytoarchitectures and activity states (Figure 1C).

To isolate individual neuron activity, we applied constrained nonnegative matrix factorization (CNMF) [24, 25], which extracts spatially segmented neurons and their calcium activity traces from each movie, giving a neuron-by-time activity matrix. The population activity of the BO slice in Figure 1A is shown in Figure 1B, where population-wide bursts are visible. Neurons differ widely in how much their activity fluctuates, so we standardized each trace to keep a few loud neurons from dominating the population covariance matrix. To capture covariation across time lags, we delay-embedded each trace, representing every time point by the current activity frame together with several past frames concatenated into one larger vector. Principal component analysis of this representation then extracts the dominant patterns of covariation (Figure 1C; see Methods IV C) [26]. We call the resulting patterns dynamical principal components, or dynamical components for short, each describing how the activity of neurons is coordinated over time. However, finite recording duration, autocorrelation, and the delay-embedding structure can produce a non-zero covariance even without correlated population activity [27]. We therefore compared the empirical component-variance spectrum with a matched surrogate null model that preserves these effects (see Methods IV D). This null model gives an upper cutoff for the largest component variance expected without coordinated population structure, and we call dynamical components with variance above that cutoff *significant* in the analysis below (see Appendix Figure 5A,D).

**FIG. 5.**
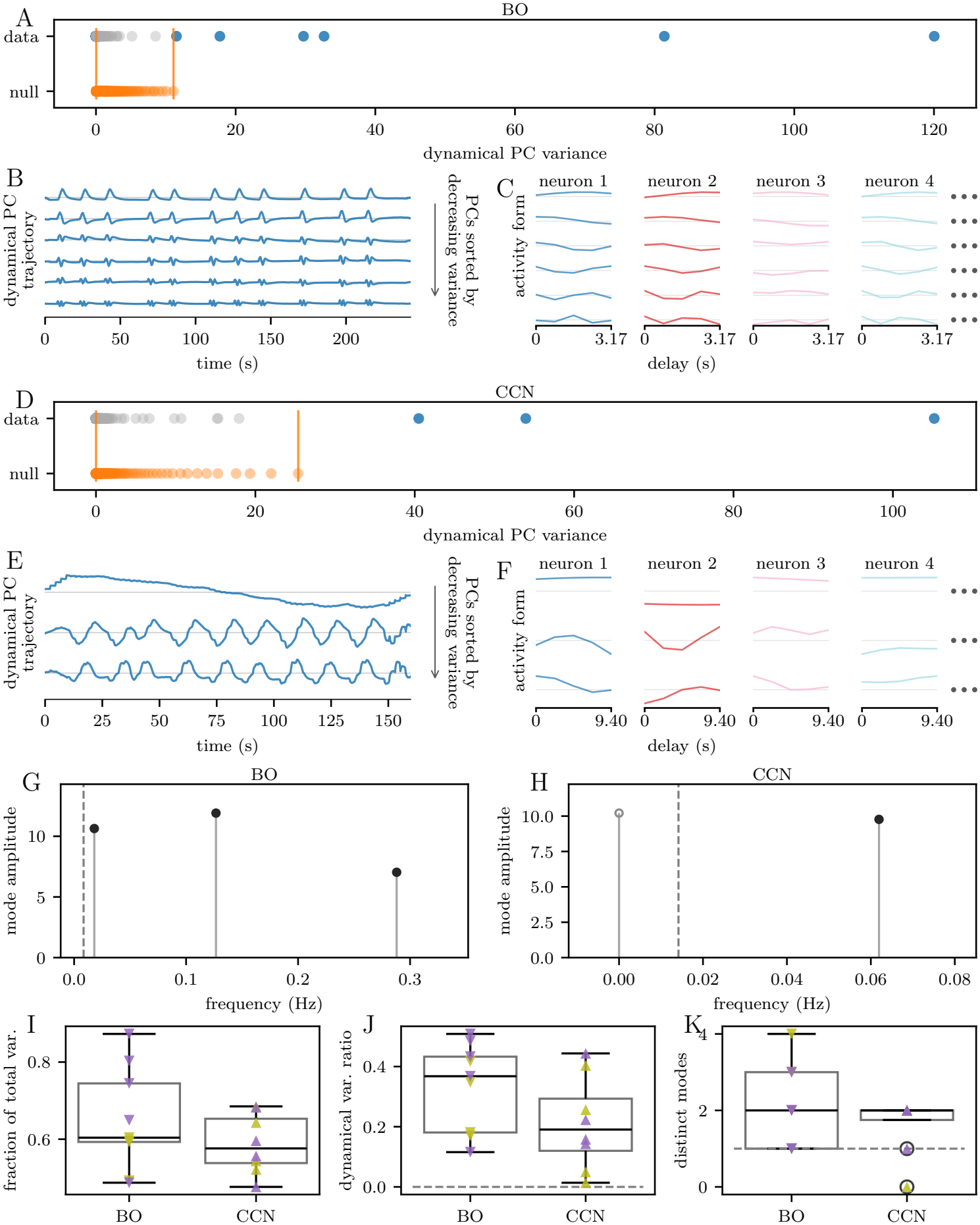
Significant Dynamical Principal Components (PCs) Account for Most of the Variance in Both Cytoarchitectures. A–F: One BO example recording under bicuculline (A–C), also shown in Figure 1A and B, and one pharmacologically excited CCN example recording (D–F). A,D: Dynamical PC variance spectra of the data and of the surrogate null. Orange vertical lines delimit the surrogate range, the dynamical PC variances expected from the null; blue markers denote significant dynamical PCs beyond that range and gray markers denote dynamical PCs within it. B,E: Trajectories of the significant dynamical PCs, sorted by decreasing variance; small step-like fluctuations at the beginning and end reflect delay-embedding boundary effects and are not interpreted as dynamics. Lower-variance dynamical PCs vary on faster timescales. C,F: Activity of example neurons within each significant dynamical PC over the delay window, with neurons arranged in columns and dynamical PCs arranged in rows by decreasing variance. Faint gray lines mark zero activity. Dynamical PCs differ in temporal variation and neuronal synchrony, with lower-variance dynamical PCs showing faster variation within the delay window. G,H: Amplitude against frequency for the dynamical modes of the fitted propagator, one point per conjugate pair or real eigenvalue, for the BO (G) and CCN (H) recording. Amplitude is the root-mean-square of a dynamical mode over the recording; the dashed vertical line marks the two-cycle resolution floor, and gray markers fall below it. The dynamical PCs entering each fit differ severalfold in variance while the dynamical modes emerge with comparable amplitudes. I: Fraction of total variance outside the surrogate range for BOs and CCNs. J: Dynamical variance ratio, where larger values indicate more power in significant dynamical PCs varying over the delay window. K: Number of dynamically distinct modes resolved by the fitted propagator, merging harmonics of a common fundamental. J,K: Dashed horizontal lines mark the reference values zero and one. None of I–K distinguishes the two cytoarchitectures. Statistics: I,J,K, unpaired Welch t-tests comparing BOs and CCNs; J,K, Holm-corrected one-sample one-sided t-tests against the reference value.

The surviving components indeed captured burst activity, varying most strongly around network bursts and spanning different timescales in the example recordings (Appendix Figure 5B,E). Across recordings, the significant dynamical components accounted for most of the embedded variance, typically between 50% and 80% (Figure 1E; Appendix Figure 5I). Figure 1E also shows that the variance spectra of the significant dynamical components were nearly the same for the BO and the CCN examples. In what follows we focus only on the activity attributed to the significant dynamical components, which we call the *denoised* activity (detailed in Methods IV E). The denoised activity preserves coordinated population bursts and removes neuron-specific fluctuations, as illustrated, for example, by the disappearance of the conspicuous orange trace in Figure 1D.

The first question we asked, i.e., whether bursts contain one activity pattern or several, can now be answered by estimating the dimensionality of the denoised activity, that is, the number of independent patterns it contains. A naive way of characterizing this is to simply count the significant dynamical components directly. However, this approach runs the risk of treating both a large dynamical component and one that only just exceeds the null model-based cutoff as equivalent, and so would classify a system with many barely significant components as high-dimensional. To address the distinct contributions of different components, we thus computed the effective dimensionality of every recording via the participation ratio [28], which weights significant dynamical components by their variance (see Methods IV D). Figure 1F summarizes these values across recordings. Our *main finding* is that, in both cytoarchitectures, the effective dimensionality was significantly greater than one (BO: *p*_Holm_ = 1.7 ×10^−4^; CCN: *p*_Holm_ = 2.0 ×10^−4^), which shows that the repertoire is multidimensional even after denoising and indicates substantial dynamical diversity. The existence of the multidimensionality thus does not depend on a particular protocol-controlled macroscopic tissue structure. Nonetheless, BOs supported a larger effective dimensionality of dynamics than CCNs (3.89 to 2.18; Welch test, *p* = 0.0043), consistent with the expectation that richer cytoarchitecture supports richer dynamics.

Several collective patterns may coexist without participating in a common dynamical process, so we asked whether the significant components evolve together, with definite timescales. We therefore regressed the components at each time point on those at the previous one and identified the eigenvectors and eigenvalues of the matrix of prediction coefficients, the *propagator* (Figure 1C, step 3; Methods IV F). The method is known as *dynamic mode decomposition* [26]. The identified eigenvectors are the linearly independently evolving *dynamical modes*, while the modulus and argument of the corresponding eigenvalue determine the growth rate and oscillation frequency of the mode. Across recordings, we typically resolved two dynamically distinct modes (Appendix Figure 5K). Despite severalfold differences in variance among the significant dynamical components that contribute to the modes, the dynamical modes emerge with comparable amplitudes (Appendix Figure 5G,H), so the dynamics are distributed quite evenly. The fitted dynamical modes showed no systematic tendency to grow or decay, tested on the dimensionless eigenvalue modulus (*p* = 0.78; see Methods IV F), so we focused on their oscillation frequencies.

The lowest of these frequencies, the fundamental frequency, is the slowest oscillation present in a recording’s collective activity (Figure 1G). Two CCN recordings, each containing only two bursts, resolved no oscillation above the two-cycle floor and were omitted. Across the remainder of the recordings, the fundamental frequency was indistinguishable between cytoarchitectures (BO mean 0.092 Hz, CCN mean 0.078 Hz; Welch test, *p* = 0.70), with the two ranges overlapping across their full extent. In these preparations, that is, in the individual organoids and networks, collective activity therefore runs at roughly one cycle per ten to twenty seconds independent of the architecture, even where repertoire size varies. The particular collective patterns and dynamical modes are recording-specific, and two preparations of the same cytoarchitecture share a repertoire no more closely than two preparations of different ones. What recurs across recordings is therefore only the *organization* of the dynamics, while the particular patterns differ.

The remaining question is whether these properties depend on the state of the network. Overall activity, and burst rate in particular, varies considerably from preparation to preparation, so we tested this by deliberately perturbing the burst rates. That is, we recorded each cytoarchitecture under two conditions, a spontaneous baseline and a pharmacologically excited condition, using bicuculline in BOs and tetraethylammonium in CCNs (see Methods IV A). Each drug is known to raise population activity reliably in its preparation, and the two act by unrelated mechanisms: bicuculline reduces GABAergic inhibition, whereas tetraethylammonium blocks potassium conductance and so raises intrinsic excitability. Across all spontaneous and excited recordings, including those with no detected bursts, pharmacological excitation raised the mean burst rate from 0.68 to 2.55 bursts per minute (*n* = 14 spontaneous and 11 excited recordings; one-sided Welch test, *p* = 0.035; Appendix Figure 6C). In contrast, excitation produced no significant change in effective dimensionality, fundamental frequency, or any other measured property of the organization of denoised dynamics in the multi-burst recordings (Appendix Figure 6D–G). In other words, excitation changes how much activity the network produces, but it leaves every measure of how that activity is organized unchanged. In particular, this allows us to pool the baseline and the excited conditions when comparing cytoarchitectures in all subsequent analyses.

**FIG. 6.**
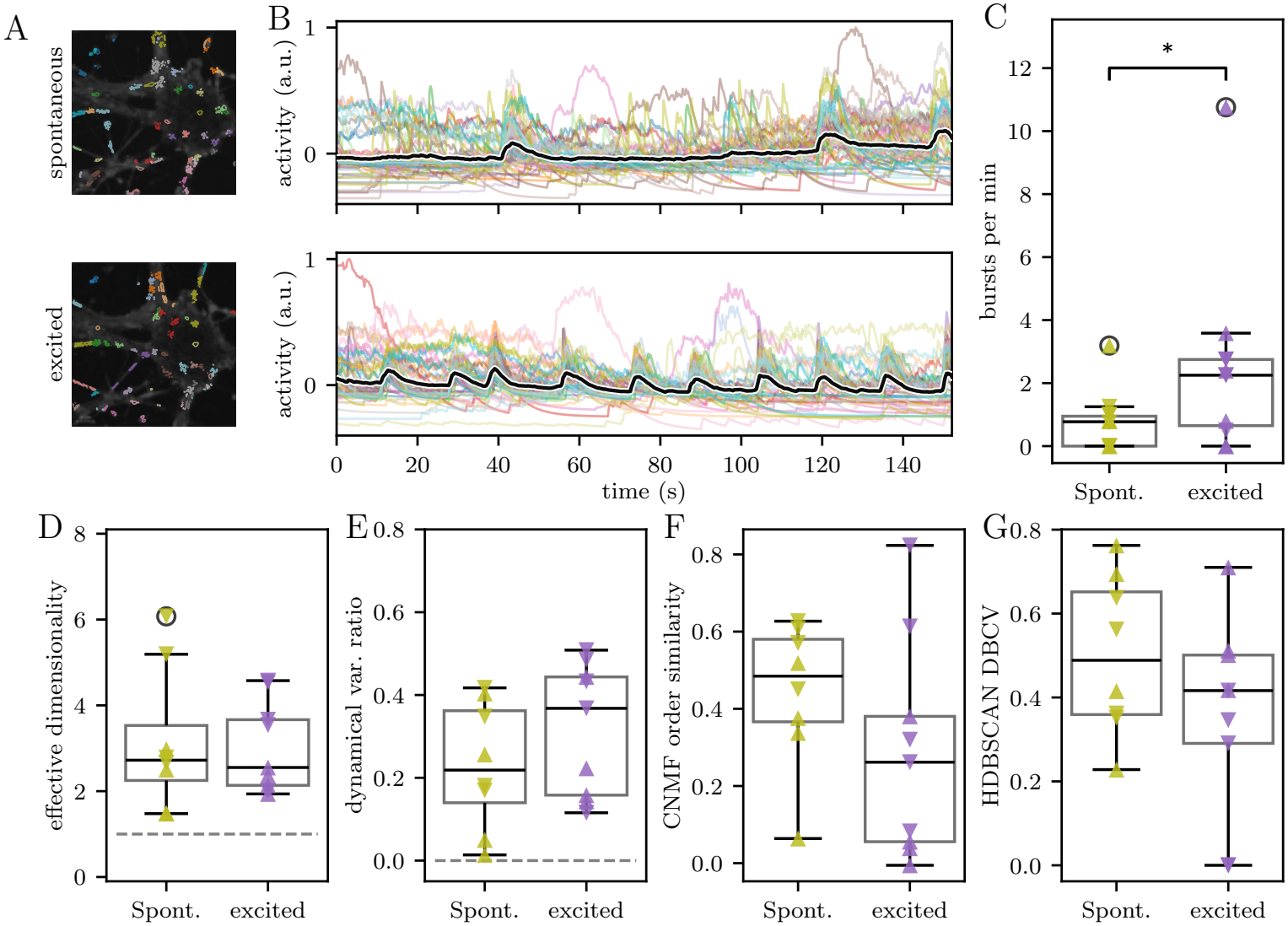
Pharmacological Excitation Increases Bursting Without Altering Other Measured Properties of Denoised Dynamics. A: CNMF segmentation of the CCN of Appendix Figure 5D–F under spontaneous and pharmacologically excited conditions. Colored contours mark identified neurons. B: CNMF activity of the same network under each condition; the black line gives mean population activity. Excitation increases the frequency of population bursts. C: Burst rate, the number of detected bursts per minute of recording, across all spontaneous and excited recordings, including recordings with no detected bursts. Excitation increases the burst rate. D–G: Effective dimensionality (D), dynamical variance ratio (E), order stability (F), and activity-form DBCV (G) under spontaneous and excited conditions among the multi-burst recordings. Excitation does not significantly change any of them. Statistics: C, one-sided unpaired Welch t-test of whether excitation increases the burst rate across all spontaneous and excited recordings; D–G, two-sided unpaired Welch t-tests comparing conditions among the 17 multi-burst recordings, with the two-way ANOVAs of Table II also run.

In summary, bursts in both cytoarchitectures are built from several distinct collective components, which combine in roughly two dynamically distinct modes, so no single pattern describes them. BOs support a larger repertoire of components, while the presence of a repertoire at all, and the properties of its dynamics, survive a change of preparation that also alters cell sourcing, differentiation age, calcium indicator, imaging medium, and even the drugs used to raise excitability (Methods IV A), one removing synaptic inhibition and the other raising intrinsic excitability. The same organization survives a severalfold pharmacological increase in overall activity, so it depends little on the network state. This stability suggests that the organization of collective dynamics may be a target of self-organization during human brain development.

### B. The Recurrent Activity Has a Graded Backbone

Bursts in both cytoarchitectures contain multiple collective patterns, more diverse than the previously reported recurrent activation sequences organized around a dedicated backbone population [14, 23]. If our results and these prior observations are both correct, the sequence participation may be graded rather than fixed: neurons could contribute to the recurrent sequence with different probabilities while remaining embedded within the same collective dynamics. We therefore asked whether recurrent activation sequences are present in our preparations and, if so, how a neuron’s contribution to the collective dynamics depends on its participation in those sequences.

To look for the recurrent order, we detected network bursts and defined an analysis window around each event (Methods IV B). Within each window, a neuron’s activation latency is its activation time relative to the other neurons (Methods IV G). When neurons were sorted by their mean latency across bursts, each burst showed a band of activity sweeping down the population in nearly the same order (Figure 2A-left). Consistent with this observation, the latency matrix, where color denotes each neuron’s latency, showed a broadly preserved activation order across burst columns (Figure 2A-right). Recurrent activation sequences are thus present in our preparations as well.

Although the sequence was readily apparent, the latency matrix also contained substantial burst-to-burst variability (Figure 2A-right). Much of this variability arises because some neurons contribute weakly or not at all to individual bursts, and the latencies assigned to them are then poorly defined. We therefore worked from the denoised traces (Methods IV E) and counted a neuron as recruited into a burst only if its activity rose more than one standard deviation above its own background (Methods IV G). The other entries of the latency matrix were blanked (Figure 2B-right; Appendix Figure 7). Once those entries were blanked, individual neurons occupied consistent positions in the sequence across bursts. Thus, variability in the sequence arises mainly from whether a neuron participates in a given burst, since its relative position when it does participate is stable, which Appendix Figure 7B quantifies over all burst pairs.

**FIG. 7.**
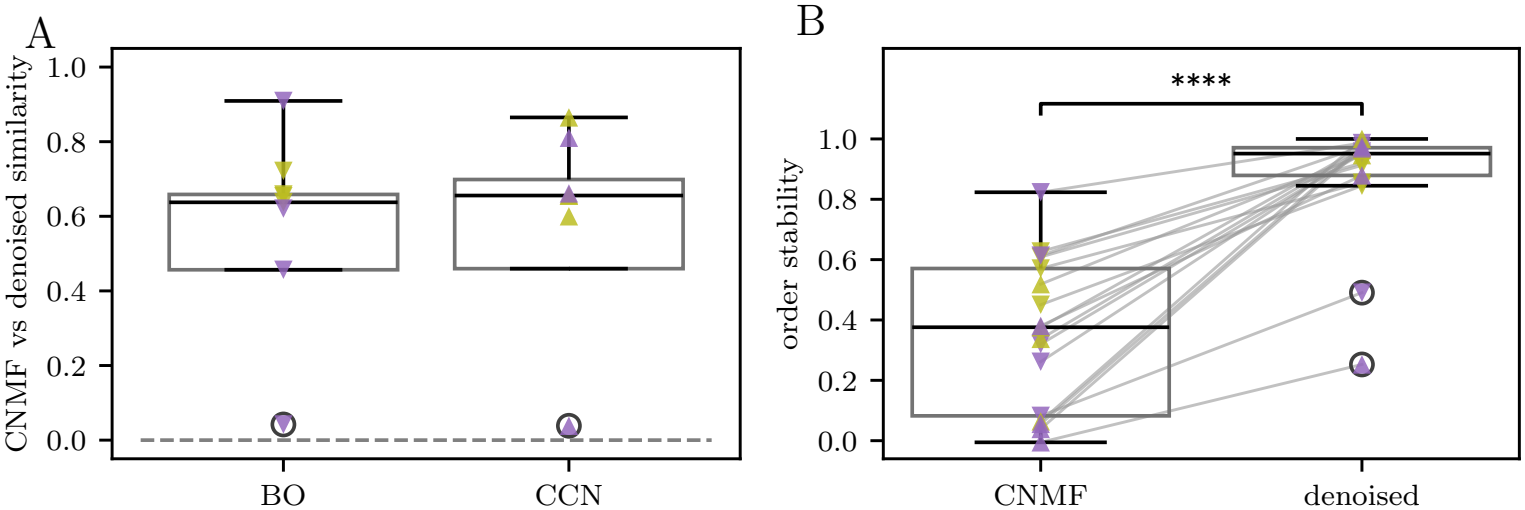
Denoising Preserves Activation Order. A: Kendall similarity between the mean activation latencies from CNMF activity and from denoised activity, one point per recording; the dashed line marks zero, and two recordings with negative similarity lie below the axis. Denoising preserves the mean activation order in most recordings. B: Order stability, the average Kendall similarity of the activation order across burst pairs, from CNMF and from denoised activity; gray lines connect the same recording. Denoised activity has the more stable order. Statistics: A, unpaired Welch t-test comparing BOs and CCNs; B, paired t-test.

The blanked latency matrix reveals a softer form of backbone organization than previously described: neurons may participate in one burst and be absent from the next while keeping a stable sequence position whenever they are recruited (Figure 2B-right). We therefore classified neurons as *consistent* if they participated in every burst, *non-participating* if in none, and *intermittent* otherwise. Intermittent neurons occurred in eight of nine BO recordings and all eight CCN recordings, accounting for 10.6% and 13.5% of the recorded populations on average (Welch test, *p* = 0.64; Figure 2C). The backbone is thus graded: participation varies from burst to burst while the recurrent order is preserved, so that several collective patterns coexist with one order without splitting neurons into a fixed backbone and a remainder.

Having established that sequence participation is graded, we next asked whether participation in the sequence predicts a neuron’s contribution to the underlying collective dynamics. To quantify this contribution, we measured the fraction of variance in the denoised population activity associated with each neuron relative to the average neuron (normalized squared loading; Methods IV H; Appendix Figure 8A). As expected, neurons that never joined the sequence contributed less strongly to the collective dynamics than neurons that participated at least intermittently (Figure 2D). Surprisingly, however, intermittent and consistent participants did not differ from one another (ANOVA, *p* = 1.0×10^−5^; non-participating vs. intermittent, *p*_Holm_ = 3.1 ×10^−4^; non-participating vs. consistent, *p*_Holm_ = 2.2×10^−4^; intermittent vs. consistent, *p*_Holm_ = 0.67). The same relationship was observed in both cytoarchitectures (participation type by cytoarchitecture interaction, *p* = 0.94). Moreover, all three participation types exhibited normalized squared loadings well above zero (one-sample tests, all *p*_Holm_ < 10^−7^), so every type contributes to the shared dynamics. Sequence participation therefore maps only coarsely onto dynamical involvement, and even neurons that never enter the sequence remain embedded in the collective dynamics (Appendix Figure 8D). Consistent with this, significant dynamical components were present even in recordings that did not exhibit bursts (Appendix Figure 8E–F), so collective dynamics extend beyond the recurrent sequence itself.

**FIG. 8.**
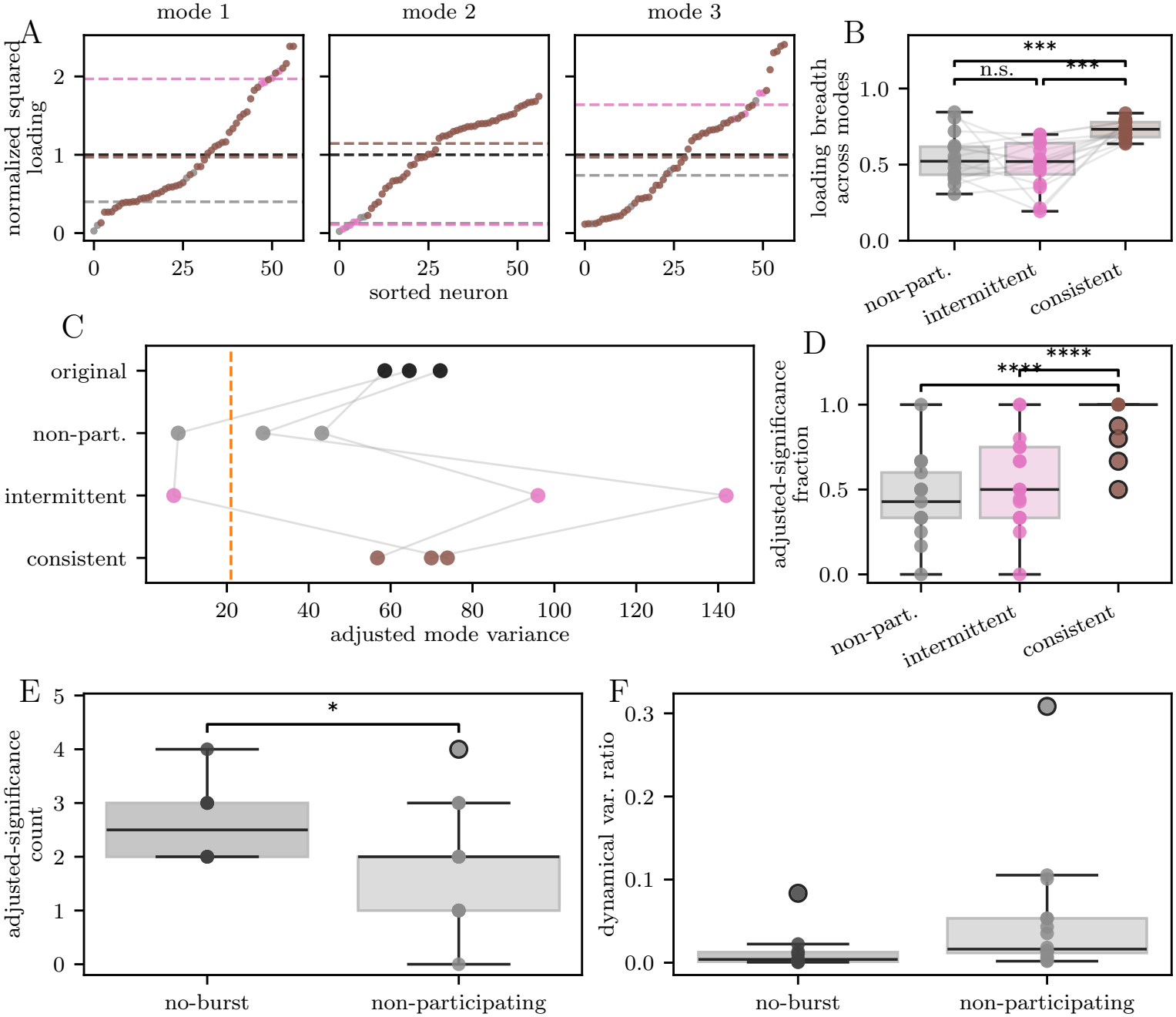
Loading of Significant Dynamical Principal Components (PCs) Extends Beyond Sequence Participants. A: Normalized squared loading of every neuron in each significant dynamical PC of the CCN recording of Figure 2, sorted within each dynamical PC and colored by participation type; black dashed lines mark the mean across all neurons, and colored dashed lines mark the mean loading of each type. B: Loading breadth, the participation ratio of a neuron’s normalized squared loadings across the significant dynamical PCs, averaged per recording within each participation type; larger values mean loading spread more evenly across dynamical PCs; gray lines connect the types within recordings containing all three. C: Dynamical PC spectrum of the recording in A, original and adjusted to the loading of each participation type; gray lines connect the same dynamical PC, and the orange dashed line marks the upper edge of the surrogate spectrum. D: Adjusted-significance fraction, the fraction of a recording’s significant dynamical PCs that stay above the edge after adjustment, for each participation type. E: Number of significant dynamical PCs in non-bursting recordings and number surviving adjustment to non-participating loading in bursting recordings. F: Dynamical variance ratio in non-bursting recordings and for non-participating neurons in bursting recordings. Statistics: B, Friedman test across participation types and Holm-corrected paired t-tests between types; D, one-way ANOVA across participation types, Holm-corrected unpaired Welch t-tests between types, and Holm-corrected one-sample one-sided t-tests against zero; E,F, unpaired Welch t-tests comparing non-bursting recordings with non-participating neurons, and Holm-corrected one-sample one-sided t-tests against zero.

Even though neurons from all participation types contribute to the collective dynamics, they may do so on different timescales. In particular, one might expect consistent sequence participants to contribute primarily to the rapid dynamics associated with bursts, whereas intermittent and non-participating neurons might contribute more strongly to slower activity. To test this possibility, we measured the dynamical variance ratio, which approaches one when neurons’ denoised activity varies substantially within individual bursts and approaches zero when it remains nearly constant over the duration of a burst (Methods IV H; Appendix Figure 5J). Consistent neurons exhibited substantially higher dynamical variance ratios than both intermittent and non-participating neurons (Figure 2E; ANOVA, *p* = 4.3 ×10^−9^; non-participating vs. consistent and intermittent vs. consistent, both *p*_Holm_ = 1.3× 10^−5^), whereas the latter two groups did not differ from one another (*p*_Holm_ = 0.72). The same pattern was observed in both cytoarchitectures, with no detectable participation-typeby-cytoarchitecture interaction (*p* = 0.37). Thus, burst-timescale fluctuations are enriched among consistent sequence participants, but are not progressively graded across all three participation types. Importantly, this distinction remains quantitative rather than categorical. The dynamical variance ratio of consistent neurons varied widely across recordings, and its values formed a continuous distribution better described by a single component than by separate populations (Appendix Figure 9). Participation type therefore predicts a neuron’s involvement in burst-timescale dynamics only on average, without partitioning the population into distinct dynamical groups.

**FIG. 9.**
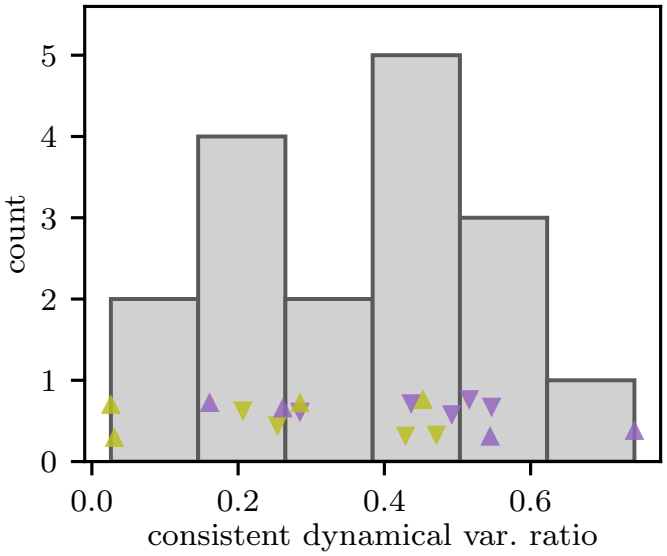
Consistent Neurons Vary Widely in the Amount of Short-Timescale Dynamics. Histogram of the dynamical variance ratio of consistent neurons, one value per multi-burst recording, with the recordings over-laid and marked by condition and cytoarchitecture. Statistics: Gaussian-mixture comparison of one-component and two-component fits by BIC.

Together, these results confirm that recurrent activation sequences are a robust feature of activity in both cytoarchitectures, but that participation in those sequences is graded rather than fixed. Neurons enter and leave the sequence across bursts, neurons that never join nevertheless contribute to the shared dynamics, and consistent participants differ from other neurons primarily in degree rather than in kind. Recurrent sequences therefore represent one manifestation of a broader collective organization that remains detectable even in neurons that rarely or never participate in bursts. Notably, the same overall organization was observed in both BOs and CCNs, with no detectable cytoarchitecture-dependent differences in participation, dynamical involvement, or burst-timescale activity. These findings suggest that key aspects of sequence organization are conserved across the distinct neural architectures examined here.

### C. Activity Types Capture Organization Beyond Recurrent Sequences

The previous section established that participation in collective dynamics is graded rather than categorical. However, participation strength alone does not fully describe a neuron’s role in the dynamics. Two neurons may contribute equally strongly while exhibiting very different activity patterns through time. We therefore asked whether neurons can be grouped according to the form of their contribution to collective activity, independent of how frequently or reliably they participate. To address this question, we characterized the normalized, delay-resolved activity profile of each neuron, which we call its *activity form*, and clustered neurons with similar forms (Methods IV I), asking also whether the resulting groups correlate with participation types.

In an example BO recording, such clustering revealed two groups separated by a clear gap (Figure 3A). To determine whether the two detected groups represent genuine structure in the data, we quantified how well separated they were by asking whether neurons were more similar to members of their own group than to members of other groups. We evaluated this separation using the density-based cluster-validity index (DBCV; Methods IV I). Across recordings, clusters were indeed significantly separated (one-sample test against zero, *p*_Holm_ = 1.8 ×10^−8^; Figure 3D), and the separation was similar in both cytoarchitectures (Figure 3C; BO mean 0.330, CCN mean 0.530; Welch test, *p* = 0.056). Two clusters per recording was the typical outcome, with a median of two in each cytoarchitecture (Appendix Figure 10C). We refer to these groups as *activity types*. The two types are distinguished primarily by a single slow component of the collective dynamics: activity in one type increases when activity in the other decreases, whereas both types show similar behavior along the remaining faster components (Appendix Figure 11A–C). Thus, neurons organize into a small number of activity types. Because both cytoarchitectures showed them, their emergence does not appear to depend on a particular large-scale tissue arrangement.

**FIG. 10.**
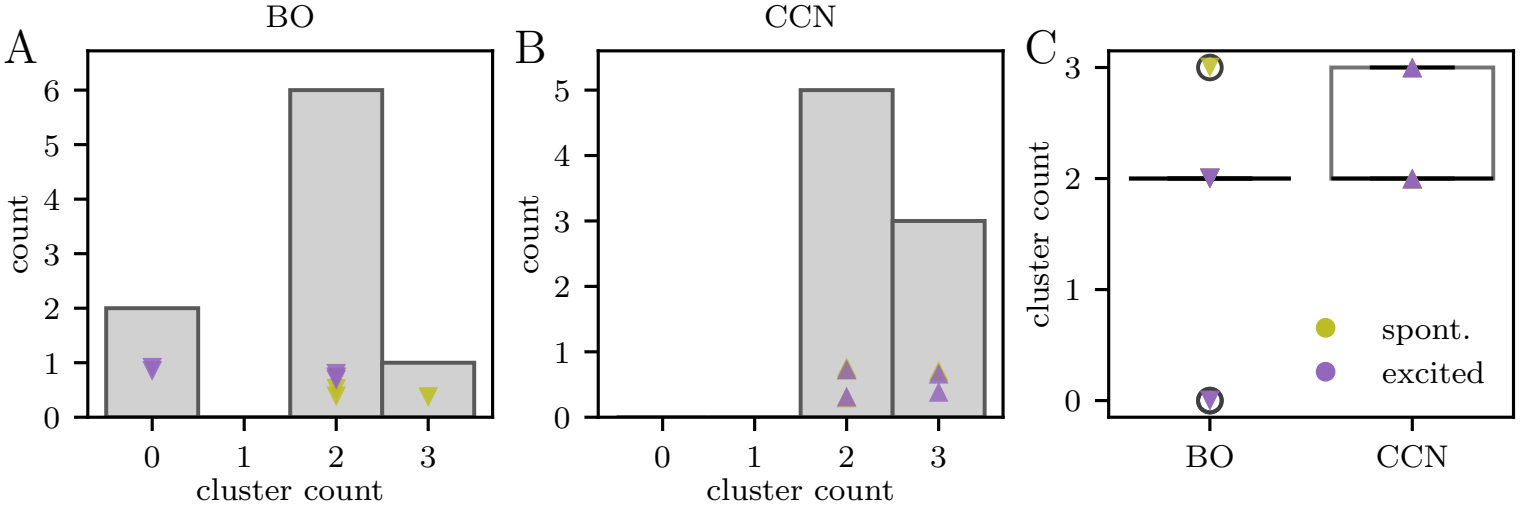
The Number of Activity Types Is Similar Across Cytoarchitectures. A,B: Distribution of the number of activity types per recording across BO (A) and CCN (B) recordings; points mark individual recordings. C: The same counts for BOs and CCNs; the number of activity types does not differ significantly between cytoarchitectures. Statistics: C, unpaired Welch t-test comparing BOs and CCNs.

**FIG. 11.**
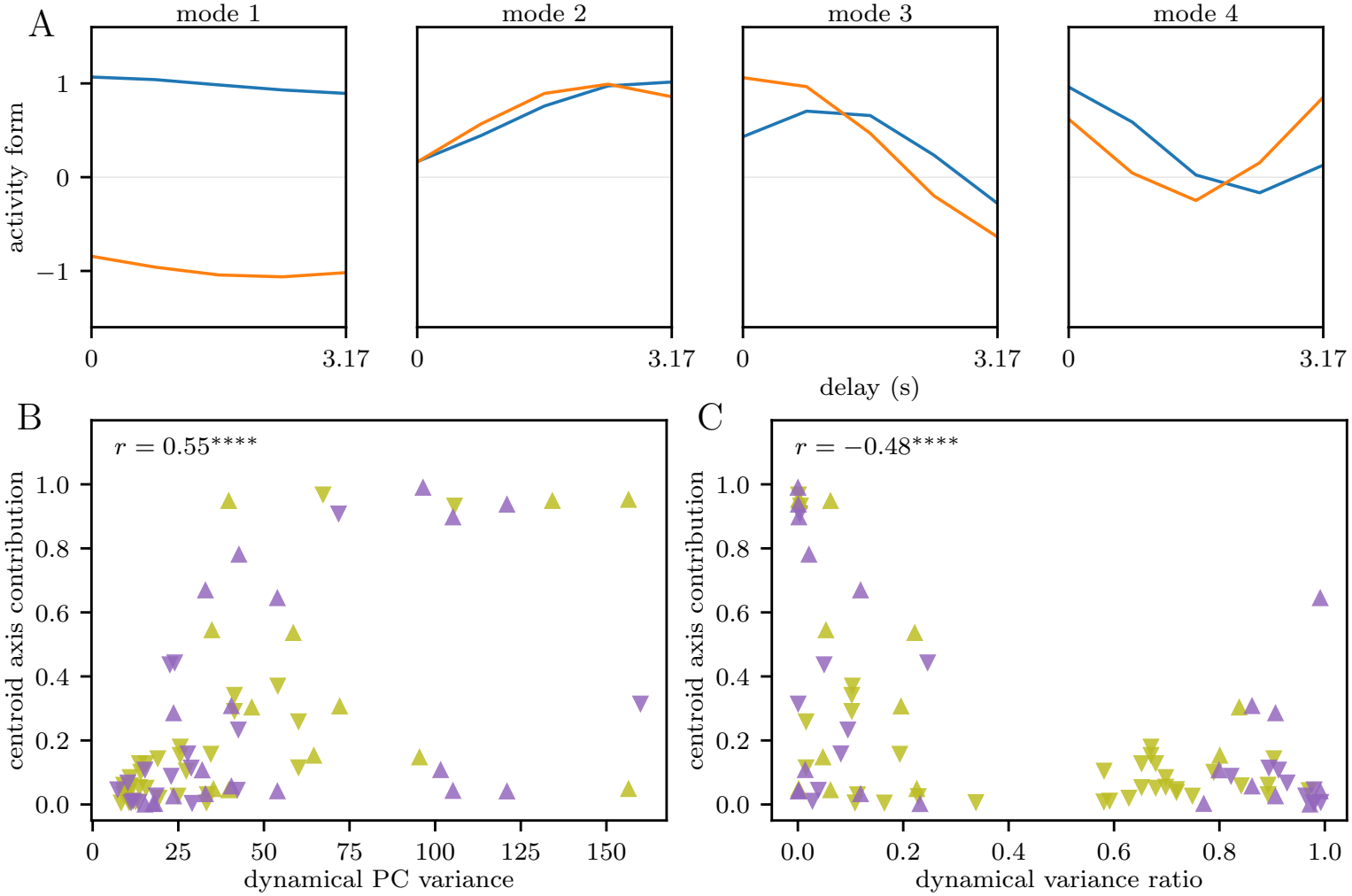
Activity Types Are Separated by High-Variance, Slow Significant Dynamical Principal Components (PCs). A: Centroid activity form of each activity type in the BO recording of Figure 3, one panel per significant dynamical PC in order of decreasing variance; each line is the mean normalized activity form of one type. The types take opposite signs on one nearly flat dynamical PC and overlap elsewhere. B: Share of a centroid axis falling on a dynamical PC vs. the variance of that dynamical PC, one point per dynamical PC, axis, and recording. Higher-variance dynamical PCs separate the types more. C: The same share vs. the dynamical variance ratio of the dynamical PC. Dynamical PCs with more short-timescale variation separate the types less. B,C: Marker shape denotes cytoarchitecture and color denotes condition, following Figure 1; text gives the Pearson correlation. Statistics: B,C, Pearson correlation.

We next asked how activity types relate to the neuronal properties identified above, particularly sequence participation and activation latency. If activity types simply reflected sequence-participation categories, neurons with the same participation type should cluster together when compared by activity-form similarity. In the example recording, however, each activity type contained a mixture of consistent, intermittent, and non-participating neurons (Figure 3A).

We quantified this across recordings by scoring participation labels with the same DBCV (Figure 3D). Whereas activity types formed well-separated groups under activity-form similarity, participation labels did not (cluster labels, mean 0.480, one-sample *p*_Holm_ = 1.8× 10^−8^; participation labels, mean −0.369, one-sample *p*_Holm_ = 0.0043; paired test, *n* = 15, *p* = 3.7 ×10^−5^). Thus, sequence participation and activity type describe distinct aspects of neuronal dynamics, partitioning the same neuronal population in different ways.

A second possibility is that activity types simply reflect a neuron’s average position within the activation sequence. Neurons activating at similar times might be expected to exhibit similar activity forms and therefore belong to the same activity type. We found that this is not the case. In the example recording, neurons spanning a broad range of activation latencies appeared within each activity type, with no obvious separation by sequence position (Figure 3B).

We tested this across recordings by representing each neuron through its activation-latency profile across bursts and asking whether neurons with similar latency profiles formed distinct clusters (Methods IV I). If activity types reflected sequence timing, recordings with strongly separated activity types should also exhibit strongly separated latency-based groups. Instead, activity-form and latency-profile DBCV scores were unrelated across recordings (Pearson *r* = −0.207, *p* = 0.43; Figure 3E). Activity types therefore capture organizational structure that is not explained by either sequence membership or sequence timing.

In summary, our results suggest that neurons organize into a small number of activity types defined by the form of their contribution to collective dynamics. These activity types are unrelated to both whether a neuron participates in the recurrent sequence and when it activates within that sequence. The same activity-type organization was observed in both cytoarchitectures, suggesting that it represents a robust feature of collective neural dynamics rather than a consequence of a particular tissue architecture.

### D. Activity Types Overlap in Space, and Activity Spreads Within Each of Them

The activity types are defined by activity alone, but it is possible that activity is determined by where neurons are located within the tissue. Similar activity often reflects shared circuitry [29], and circuitry has pronounced spatial structure, so activity types could be correlated with it in several ways. One possibility is that each activity type occupies its own spatial neighborhood, its neurons similarly active because nearby neurons are the most likely to be connected in developing cortex [30, 31]. Another is that neurons form pathways, and activity preferentially spreads through strongly connected neurons, as spontaneous waves do in developing circuits before sensory experience [1, 2]. Here we test both possibilities.

The possibility of spatial neighborhoods was not supported by the spatial distribution of neurons in the example recording, where neurons assigned to different activity types were extensively intermixed in space and occupied no separate territories (Figure 4A). Across recordings, scoring the activity-type labels with DBCV on the physical distances between neurons (Methods IV J) gave significantly negative values in both cytoarchitectures, so neurons of the same activity type are no closer to one another than to neurons of the other types (Figure 4C; BO, *p*_Holm_ = 1.1 ×10^−4^; CCN, *p*_Holm_ = 1.3 ×10^−4^). These results therefore are not consistent with the possibility that activity types represent coordinated highly localized spatial activity patterns.

We next considered the possibility of activity propagation through pathways of highly coupled neurons. If activity travels through such pathways, neurons farther from where it starts should activate later. We therefore took the earliest-activating neurons as the origin and correlated each neuron’s latency with its distance from that origin (Methods IV J). The larger the correlation, the more closely timing follows space. The example recording shown in Figure 4A exhibited little relationship between activation latency and physical distance across all recruited neurons. In contrast, significant distance– latency correlations emerged within each activity type (Figure 4B). Thus, each activity type in this recording exhibited its own spatial ordering of activation, consistent with partially independent propagation-like dynamics.

Similar findings held across the recordings: the distance–latency correlation was higher within activity types than across the whole population of the same recording (Figure 4D). This within-type increase was clear in BOs (one-sample test, *p*_Holm_ = 0.0039), while in CCNs alone it was not significant (*p*_Holm_ = 0.27); however, the two cytoarchitectures did not differ significantly (*p* = 0.16), and the increase was significantly positive when all recordings were pooled (mean 0.153, *p* = 0.012).

Together, these observations argue against the interpretation that activity types reflect simple spatial neighborhoods. Instead, neurons belonging to the same activity type are spatially intermixed yet exhibit stronger within-type spatiotemporal ordering, consistent with overlapping functional subnetworks that support partially independent propagation-like dynamics.

## III. DISCUSSION

In this study, we explored how the collective dynamics of self-organized neural tissue depend on large-scale architecture. Surprisingly, many organizational features of collective dynamics were conserved across the two architectures despite their substantial anatomical differences. Both BOs and CCNs exhibited bursts composed of multiple collective patterns, recurrent activation sequences with graded backbone participation, and a small number of activity types displaying propagation-like spatial organization. Among the properties examined here, the size of the collective repertoire was the only organizational feature that differed detectably between the two cytoarchitectures, and none of the measured organizational features changed when pharmacological excitation increased burst rates severalfold. Together, these observations suggest that several organizational principles of collective dynamics are remarkably robust across distinct self-organized neural architectures.

What determines this organization of activity? The two cytoarchitectures differ in cell sourcing, differentiation age, calcium indicator, imaging conditions, and especially in mesoscale organization, contrasting continuous tissue with spatially separated neuronal clusters connected by neurite bundles. At the same time, they share many cellular constituents and local developmental processes. Our results therefore suggest that several organizational properties of collective dynamics may arise from features common to both systems. Largescale architecture nevertheless remains important. BOs support a larger repertoire of collective patterns than CCNs, indicating that architecture modulates the richness of the dynamics. Yet many organizational features remain conserved despite markedly different mesoscale organizations. Thus, architecture appears to influence the diversity of collective activity more strongly than its basic organizational framework. Whether this principle extends beyond the architectures examined here remains an important question for future work.

Our analysis extends recent interpretations of organoid activity that emphasize fixed sequences and dedicated backbone populations in self-organized neural tissues [23]. In our data, backbone participation is graded: membership shifts from burst to burst, and even neurons that never join the sequence remain embedded within the collective dynamics. Zebra finch HVC exhibits a related dissociation between reproducible temporal order and fixed neuronal membership, although in the context of behavior, where song-locked timing remains stable across days while participation varies across renditions [32]. By contrast, theoretical constructs such as synfire chains rely on highly structured connectivity for rigid propagation through successive neuronal groups [33]. Together, these observations raise the possibility that rigid propagation requires specialized wiring, whereas self-organized connectivity may naturally give rise to the graded, multipattern organization observed here and in other neural systems.

Each of these findings has a counterpart in the spontaneous activity of the intact brain. Cortical population activity in vivo is often described by a limited number of collective activity patterns [3, 4], hippocampal and cortical circuits replay internally generated sequences [5, 6], and developing neural circuits exhibit activity waves that coordinate maturation [1, 2]. It is therefore notable that related dynamical motifs emerge in tissue models that receive no sensory input and whose fine-scale connectivity is not experimentally specified. These observations suggest that many properties of collective activity can arise robustly through developmental self-organization, even in the absence of sensory experience and experimentally designed wiring. Notably, engineered biological systems have previously enabled quantitative descriptions of collective behavior. For example, studies of cell populations in controlled environments have revealed both the benefits of collective sensing and the limits imposed by signaling and communication noise [34–36]. By analogy, engineered neural tissues may provide useful and experimentally accessible systems for investigating collective neural dynamics and their perturbation in neurodevelopmental disorders.

Our findings also have implications for how activity in engineered neural tissues is characterized. Organoids and related models are increasingly used to study development, disease mechanisms, and therapeutic responses, yet their activity is often summarized primarily through descriptors such as burst frequency or burst size. In our experiments, burst rates increased approximately fourfold under pharmacological excitation, whereas the organizational properties of collective dynamics examined here remained unchanged. This observation suggests that burst statistics alone provide only a partial description of network activity and do not fully capture the richness of the underlying collective dynamics. By contrast, the collective repertoire, graded sequence participation, dynamical modes, and activity types emerge directly from neuron-level recordings and describe aspects of population organization that conventional summary measures do not. As engineered neural tissues become increasingly important experimental systems, these dynamical properties may complement traditional activity metrics and provide a richer description of circuit organization.

Among these properties, activity types remain the least understood. We identified several distinct activity types across both cytoarchitectures, each associated with partially independent propagation-like dynamics.

Their biological basis remains unclear. They may reflect differences in cellular identity, developmental state, or functional circuit organization, each of which generates distinct experimental predictions. Combining functional imaging with molecular characterization will therefore be an important next step in determining how these activity types relate to underlying neural biology. Such experiments could clarify whether activity types arise primarily from differences in cellular identity, from circuit organization, or from interactions between the two.

Our study has several limitations. Although the dataset is sufficient to resolve the major dynamical features reported here, the number of recordings remains modest and may limit sensitivity to subtler architecturedependent effects. Recording durations are also constrained, reducing statistical power for detecting weaker forms of collective structure. In addition, the null model was estimated from detrended activity, which effectively treats slow drift as signal. Alternative treatments of lowfrequency fluctuations could therefore alter the significance threshold for weak activity components. However, we do not expect the principal conclusions to depend strongly on this choice, because all downstream analyses were dominated by the strongest collective components (Appendix Figure 11).

In summary, our results suggest that self-organization may shape not only the cellular composition and anatomy of neural tissue, but also the organization of its collective dynamics. Determining how broadly this principle generalizes will require comparable analyses across a wider range of neural architectures and developmental contexts. Complementary theoretical work will be needed to identify the structural constraints capable of generating such dynamics [37], an increasingly tractable problem given recent advances in the study of disordered neural networks [38, 39]. Finally, spatially controlled perturbations may reveal whether the activity types identified here correspond to causally connected pathways rather than statistical dynamical groupings. More broadly, collective activity patterns, graded participation, activity types, and propagation-like organization may provide a useful vocabulary for describing neural circuits that complements traditional measures such as firing rates, burst statistics, and synchrony.

## IV. METHODS

### A. Organoid Protocols and Calcium Imaging

a. *Human iPSC culture and neural induction* Human induced pluripotent stem cells (hiPSCs), reprogrammed from human fibroblasts, were maintained under standard pluripotent stem cell conditions and differentiated toward a neural lineage with a protocol based on embryoid bodies (EBs), modified from Mariani *et al*. [16], Kim *et al*. [40]. For CCN neural induction, hiPSC cultures were pretreated with ROCK inhibitor (Y27632, 5 *µ*M) for 1 h to enhance survival. Cells were dissociated with Accutase diluted 1:2 in DPBS into single-cell suspensions. Cells were seeded into AggreWell plates at 3 ×10^6^ cells per well in neural basal (NB) medium supplemented with ROCK inhibitor and recombinant Noggin (200 ng*/*mL) to promote EB formation. EBs were cultured for 2 days and then transferred to suspension culture in NB medium with Noggin. At day 4, EBs were plated onto Matrigelcoated dishes to allow attachment and radial outgrowth. Neural induction was continued in NB medium supplemented with FGF2 (20 ng*/*mL), Noggin (200 ng*/*mL), and Dkk1 (200 ng*/*mL), with medium changes every other day.Neural rosette structures were manually isolated at day 8 and replated in suspension culture with FGF2 and EGF (10 ng*/*mL each) for expansion. After approximately 1 month, the resulting neural progenitor cells (NPCs) were dissociated into single cells for downstream neuronal network formation.
b. *BO protocol* Cortical-like BOs (derived from the 5D culture area after DuoMAPS patterning) were generated and sliced as described by Scuderi *et al*. [14], and the recordings analyzed here were acquired in the course of that study. Slices prepared on terminal differentiation day 120 were seeded on poly-L-ornithine and laminin-coated plates and infected with the GCaMP8m virus at a dilution of 1:100. Cultures were incubated with the virus for 10 days. Fresh medium was added after 96 h and then every 2 days until day 10, followed by a full medium change on day 11. BOs were allowed 2–3 weeks for indicator expression before imaging at day 150 in BrainPhys Imaging Optimized Neuronal Medium (STEMCELL Technologies).
c. *CCN protocol* Neural progenitor cells (NPCs) derived from hiPSCs were used to generate large-scale self-organizing neuronal networks. Cells were dissociated and plated at high density (approximately 3×10^6^ cells*/*cm^2^) onto culture plates coated with poly-Lornithine (overnight) followed by laminin (10 *µ*g*/*mL, 2 h). Cells were initially maintained in NPC medium for 48 h to allow attachment and recovery. Medium was then switched to neuronal differentiation medium consisting of Neurobasal (Gibco) supplemented with B27 (5%), Gluta-MAX (1%), and neurotrophic factors (BDNF and GDNF at 20 ng*/*mL each). Cultures were maintained at 37 ^*°*^C with 5% CO_2_ through terminal differentiation, approximately 30 days. During this period, cells spontaneously self-organized into clustered neuronal assemblies interconnected by extensive neurite bundles, forming large-scale (approximately 1 cm^2^) functional networks, and matured progressively, with (a) emergence of heterogeneous neuronal populations, including excitatory and inhibitory neurons, (b) formation of long-range neurite bundles connecting cellular clusters, and (c) development of spontaneous electrical activity and synchronized network dynamics [19, 20].
d. *Calcium imaging under pharmacological conditions* BO slice calcium imaging was performed in BrainPhys medium, as described previously [14]. Briefly, organoid slices were imaged on differentiation day 150 using the genetically encoded indicator construct (pGP-AAV-syn-jGCaMP8m-WPRE). Recordings were acquired through a ×25, 0.8-NA water-immersion objective, bidirectionally at 512 ×512 pixels with line average 8, at one frame every 264 ms (3.78 Hz), with approximately 909 frames per recording. For BO slices, pharmacologically excited recordings used bicuculline (50 *µ*M), a GABA_*A*_ receptor antagonist. Some BO recordings also included muscimol (25 *µ*M), a GABA_*A*_ receptor agonist, and washout conditions. The muscimol recordings contained no bursts and so fell outside the multi-burst set analyzed here, while one post-muscimol washout recording was pooled with the spontaneous BO group, which therefore comprised four recordings, one of which followed a washout and was not a naive baseline. The same fields of view were re-imaged after drug application to evaluate changes in network activity. For CCN recordings, cultures at terminal differentiation day 30–45 (TD30–45) were incubated with Fluo-4 AM (2 *µ*M) for 1 h. CCN imaging was performed using confocal microscopy (488 nm excitation; approximately 261 ms per frame). CCN recordings were conducted in artificial cerebrospinal fluid (aCSF: 130 mM NaCl, 3 mM KCl, 1.25 mM NaH_2_PO_4_, 26 mM NaHCO_3_, 1 mM MgCl_2_, 1 mM CaCl_2_, 5 mM glucose). For CCNs, activity was increased by acute addition of 10 mM tetraethylammonium, and the same fields of view were re-imaged after drug application to evaluate changes in network activity. Thus, the pharmacologically excited condition was bicuculline for BOs and tetraethylammonium for CCNs.
e. *Recordings and units of analysis* Recordings were obtained from six BOs and six CCNs, each imaged under more than one pharmacological condition, giving 28 recordings in total. The 17 analyzed recordings were those containing more than one network burst: nine BO recordings from six organoids and eight CCN recordings from four networks. Each CCN contributed one spontaneous and one excited recording, while BOs contributed one to three recordings each, so the BO recordings were distributed unevenly across organoids, and the spontaneous and excited BO groups were not drawn from the same set of organoids. Ten recordings contained no bursts, comprising the spontaneous recordings of three BOs, the muscimol recordings of three others, and both conditions of the two remaining CCNs; these entered only the non-bursting comparison of Appendix Figure 8E– F. One further BO recording contained a single burst and was excluded from analyses that compared bursts within a recording. Comparisons between cytoarchitectures therefore rest on six BOs and four CCNs, but we did not treat repeated recordings of one preparation as repeated measurements, for the reasons given alongside Appendix Table I. Points in summary plots denote individual recordings.

**TABLE I.** No Preparation-Level Effect Is Detected in Repeated Recordings. Within SD is the pooled standard deviation across recordings of the same preparation, computed over preparations contributing more than one recording. Between SD is the standard deviation of preparation means. *F* and *p* are from a one-way analysis of variance with preparation as the grouping factor, testing for a preparation-level effect; no test reaches significance. Six BOs contribute nine recordings, only two BOs more than one; four CCNs contribute two recordings each.

|  | within SD | between SD | *F* | *p* |
|---|---|---|---|---|
| effective dimensionality |  |  |  |  |
| BO | 1.08 | 1.09 | 2.26 | 0.27 |
| CCN | 0.51 | 0.39 | 1.17 | 0.42 |
| dynamical variance ratio |  |  |  |  |
| BO | 0.15 | 0.15 | 1.48 | 0.40 |
| CCN | 0.16 | 0.10 | 0.73 | 0.59 |
| activity-form DBCV |  |  |  |  |
| BO | 0.15 | 0.25 | 4.37 | 0.13 |
| CCN | 0.19 | 0.12 | 0.72 | 0.59 |

### B. Burst Detection

Bursts were detected from the population mean of the standardized CNMF activity after subtracting its running median, computed over a window spanning 60% of the recording, which flattens the slowly varying baseline so that peak heights and widths were measured against a constant reference. Local maxima of the residual counted as burst peaks only if they exceeded one third of the residual’s largest value in the recording, a threshold chosen empirically. Any two peaks closer together than the width of the widest peak, measured at 60% of its prominence below the top, were treated as one burst, represented by the higher of the two, so that a single burst with two local maxima in the population mean was not counted twice. Burst windows were set to have one common width per recording, chosen so that the window fit the broadest burst: the width of the widest peak measured near its base, at 80% of its prominence below the top. When bursts followed each other closely, this width was capped at 0.75 of the smallest interval between consecutive peaks, so that the windows of neighboring bursts overlapped little. Each burst’s window then extended 0.6 of this width before its peak and 1.2 after, because bursts rise quickly and decay slowly. All detected bursts were inspected visually to confirm that they correspond to population events.

### C. Delay Embedding

The number of delay steps must balance two effects: more delays increase the dimensionality of the embedded space, which allows more expressive dynamical principal components (PCs), but also require more time points to resolve those dynamical PCs well. Based on the size and duration of the recordings, we used five delay steps. The delay spacing was set from each recording’s own autocorrelation. Each neuron’s activity was first detrended by subtracting a running median over 30% of the recording, smoothed with a cubic polynomial, so that slow drift did not inflate the autocorrelation. The mean autocorrelation over neurons was then fitted over the first five lags by an exponential, and the delay spacing was three times the lag at which this fit fell to 0.85, rounded to the nearest whole frame. Since an exponential falls to 0.85 after about one sixth of its decay time, the spacing is about half an autocorrelation time, and the five delays span four spacings, approximately two autocorrelation times, long enough to capture spatiotemporal structure within bursts while avoiding dynamical PCs dominated by slow drift. Shorter spacings were avoided because single-neuron autocorrelation would then dominate the covariance and swamp the cross-neuron structure the embedding is meant to capture.

Let *Ã*_*i,t*_ denote the standardized activity of neuron *i* at frame *t*, with zero mean and unit variance across time. This standardization, applied before embedding, keeps dynamical PC identification from being dominated by the neurons with the highest single-unit variance. The delay-embedded matrix is defined by

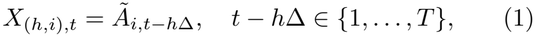

where *h* ∈ {0, …, *H* − 1} indexes the delay channel, *i* ∈ {1, …, *N*} indexes the neuron, and *T* is the recording length. Because the recording is finite, delay channel *h* is defined only for *t* ∈ {*h*Δ + 1, …, *T* + *h*Δ}, producing uneven edges: earlier channels extend to the left and later channels to the right of the original recording window. Entries outside the valid range were set to zero, so the full embedded matrix has *HN* rows and *T* + (*H* − 1)Δ columns and every original frame contributes to all *H* channels. The embedded matrix was normalized by 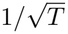 before the principal component analysis (PCA). The larger factor 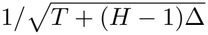 would overcount, because each row of the embedded matrix contains exactly *T* nonzero entries, so the zero-padded entries contribute no signal and the effective sample size is *T*. Burst windows were only retained if they fell entirely within {1, …, *T*}.

### D. Identification of Significant Dynamical PCs

PCA was applied to the standardized delay-embedded activity, decomposing variance into dynamical PCs ordered by their variance. Writing the decomposition as a singular-value factorization fixes the notation used throughout the Methods,

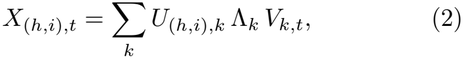

where *U*_*·,k*_ is the loading vector of dynamical PC *k*, Λ_*k*_ its singular value, and *V*_*k,·*_ its unit-norm time course. The dynamical PC variance is 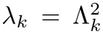. Because the dynamic mode decomposition and the reconstruction described below both act on time courses that retain the dynamical PC’s amplitude, we write that scaled combination as

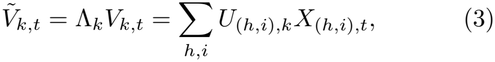

the trajectory of dynamical PC *k*. Since finite recording length, single-neuron autocorrelation, and the delay embedding itself produce a spectrum of nonzero variances even for neurons that are entirely independent of one another [27, 41], not every dynamical PC is meaningful. To tell collective structure from this background, we built a surrogate null, an ensemble of independent neurons with the same autocorrelation as the detrended data, the same recording length, and the same embedding, and we retained as significant the dynamical PCs whose variance exceeded the largest variance the null produced.

Specifically, the autocorrelation was modeled as exponential decay *r*_*τ*_ = *a*^*τ*^ , with *a* ∈ (0, 1) from the exponential fit over the first five lags described above. Null trajectories were drawn from an autoregressive process of order one (AR(1)) [42] with parameter *a*:

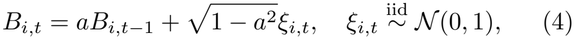

which gives ⟨*B*_*i,t*_*B*_*i,t*+*τ*_ ⟩_*ξ*_ = *a*^*τ*^ by construction, matching the fitted autocorrelation exactly. Null trajectories were then standardized and delay-embedded with the same *H*, Δ, and zero-padding as the data, so the null spectrum inherits the same embedding structure. Twenty independent null draws were generated, and the median variance across draws was taken at each dynamical PC rank, giving a single null spectrum. The spectral edge is the largest variance in that spectrum, which is its highestranked dynamical PC.

The richness of the repertoire of significant dynamical PCs was summarized by the effective dimensionality, defined as their number multiplied by the participation ratio of their variances [28], a standard measure of the dimensionality of neural population activity [38, 39, 43, 44]. The participation ratio is 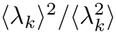, where *λ*_*k*_ are the significant dynamical PC variances, and angle brackets denote the mean over significant dynamical PCs. The ratio downweights spectra dominated by a single highvariance dynamical PC.

### E. Denoised Activity

The denoised activity is the part of each neuron’s activity explained by the significant dynamical PCs. Projecting the embedded activity onto those dynamical PCs keeps, in every delay channel, the activity that covaries with the rest of the population along the significant dynamical PCs, and suppresses the noise in the dynamical components below the significance threshold. The perneuron denoised trajectory is the average of the *H* delaychannel contributions, each shifted back to zero lag.

Concretely, in the notation of Methods IV D, the reconstruction keeps only the *K* significant dynamical PCs and their trajectories 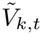. The projected embedded matrix is

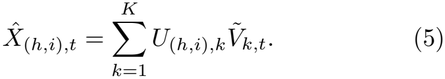

The per-neuron denoised trajectory *Â* _*i,t*_ was obtained by reversing the delay embedding: each delay channel was shifted back to zero lag and the *H* channels were averaged,

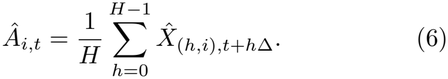

The result was then smoothed with a moving average one delay spacing Δ wide to suppress high-frequency fluctuations that can arise from the spaced delay embedding and affect local peak detection.

### F. Dynamic Mode Decomposition

We are interested in the dynamics of the significant dynamical PCs, meaning how the population state at one time determines the state at the next and how the components drive one another as the state evolves. That requires a map forward in time. PCA gives no such map. We used the simplest such map, a linear one: we fit a propagator by regressing the trajectories of the significant dynamical PCs at each frame on those at the previous frame, the method known as dynamic mode decomposition [26, 45, 46]. The fit was on 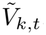, the trajectory of significant dynamical PC *k* defined above, so that dynamical-mode amplitudes inherit the dynamical PC variances. Let *T* be the set of frames at which every delay channel is filled, which excludes the zero-padded edges of the embedding. The propagator *A*, a *K* ×*K* matrix, is the least-squares solution of 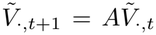 over *t*∈ *T* , fitted on the whole recording. An eigenvector *ϕ*_*j*_ of *A* is a dynamical mode in the reduced dynamical PC space, and *U*_*K*_*ϕ*_*j*_ gives its corresponding pattern in the delay-embedded neural space, where *U*_*K*_ contains the loading vectors of the *K* significant dynamical PCs. We fit over the whole recording, since a fit confined to burst windows returns frequencies close to the inverse window length, because a segment of length *L* looks like a single cycle of period *L*, and such frequencies would measure the burst detection and not the dynamics.

Noise in the current state is uncorrelated with the next state, so it dilutes the fitted dependence. This is the usual attenuation of a regression slope by errors in the predictor, and it makes every mode of a least-squares *A* look more damped than it is. *A* was therefore estimated in forward-backward form [47], 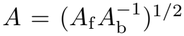, where *A*_f_ and *A*_b_ are the forward and backward least-squares solutions and the two biases, of opposite sign, cancel to first order. Writing an eigenvalue of *A* as *µ* = |*µ*| *e*^*iθ*^, the dynamical mode it labels has growth rate ln |*µ*| */δt* and oscillation frequency *θ/*(2*πδt*), for the frame interval *δt*, taken as 0.264 s for BOs and 0.261 s for CCNs. Because *A* is real, complex eigenvalues occur in conjugate pairs, and each pair contributes one frequency. A frequency is called resolved when it completes at least two cycles within the fitted frames *T* , and the lowest resolved frequency of a recording was reported as its fundamental.

Refitting the propagator at frame lags of two and three tested whether the linear description holds across timescales: since a propagator fitted at lag *L* is the lag-one propagator raised to the *L*th power, dynamical modes were matched across lags by eigenvector overlap and their frequencies compared, excluding lags at which a mode would exceed the Nyquist limit. The fitted dynamical modes were also examined for a tendency to grow or decay, through the eigenvalue modulus |*µ*| , which is dimensionless and equals one for a neutrally stable mode. Each real eigenvalue and each complexconjugate pair contributed one value, and the median was taken across dynamical modes within each of the 17 multi-burst recordings so that recordings with more significant dynamical PCs did not receive more weight. The recording-level medians were compared with one using a two-sided one-sample Wilcoxon signed-rank test. Absolute growth rates were not compared across recordings, because the delay spacing Δ was set per recording from its own autocorrelation, so quantities carrying units of time inherit that adaptation.

### G. Activation Latency, Participation, and Order Analysis

Activation latency *τ*_*i,b*_ was estimated for both CNMF and denoised activity using the same softmax-weighted center-of-mass approach. For each neuron-burst pair, a linear baseline *Ā*_*i,t*_ connecting the first and last frames of the burst *A*_*i,t*_ was subtracted to give the detrended trajectory *Ã*_*i,t*_ = *A*_*i,t*_ −*Ā*_*i,t*_, which was used to identify the support interval [*t*_1_, *t*_2_] of the largest detected peak, from the trough before it to the trough after it. Activation latency is the softmax-weighted center of the detrended activity over that interval:

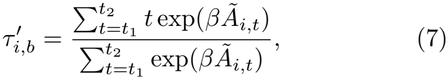

where *β* = 3 controls how sharply weight is concentrated near the peak, and the *β*→ ∞ limit recovers the argmax. Using the detrended activity focuses the estimate on short-timescale activation events and keeps slow global drift out of it. Results were not sensitive to moderate changes in *β*. Activation latency is defined by mediancentering 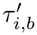 within each burst, 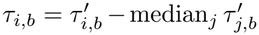, so that *τ*_*i,b*_ measures where a neuron sits in the activation order and carries no information about when the burst occurred. For CNMF activity, the traces were first smoothed with a moving average spanning the full delay window, *H*Δ, to reduce spurious peaks from noise.

For denoised activity *Â*_*i,t*_, no latency was assigned to a neuron in a burst if the maximum deviation of the detrended trajectory did not exceed a threshold set by that neuron’s residual fluctuation, or if no peak was detected. More concretely, we define the detrended denoised activity as 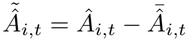, where 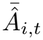 is the linear baseline of the denoised activity *Â*_*i,t*_. The threshold for neuron *i* is the standard deviation over time of its residual fluctuation *A*_*i,t*_ −*Â*_*i,t*_, the part of standardized CNMF activity *A*_*i,t*_ not captured by the *K* significant dynamical PCs. Because the threshold is defined by the residual of the denoising, no such criterion exists for raw CNMF traces, which were assigned a latency unless no peak was found.

For the burst-aligned displays, neurons were ordered by their mean CNMF activation latency, averaged over the bursts in which that neuron had an assigned latency, and the same order was used before and after denoising (Figure 2A and B).

Sequence participation for each neuron is its fraction of bursts with an assigned denoised latency. Neurons with participation near one are termed consistent, those near zero non-participating, and those in between intermittent, a graded version of the backbone criterion of participation in every burst [23]. For each of the CNMF and denoised activity representations, neurons with an assigned latency in at least one burst were eligible for the order analysis. Burst-to-burst order stability is the mean Kendall rank correlation [48] of activation latencies across all pairs of bursts. For each burst pair, the Kendall correlation was computed only over neurons with assigned latencies in both bursts; correlations from pairs with fewer than two such neurons were undefined and omitted from the recording-level mean. For the order-preservation validation in Appendix Figure 7A, the Kendall rank correlation was computed between the vector of mean CNMF latencies and the vector of mean denoised latencies across neurons.

### H. Normalized Squared Loading and Dynamical Variance Ratio

Normalized squared loading measures how much a neuron contributes to the denoised collective dynamics. It is the second moment of the neuron’s activity profile within a dynamical PC, normalized so that the across-neuron mean within each dynamical PC is one:

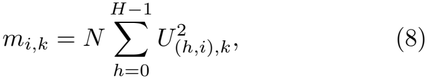

where *N* is the number of neurons in the recording, a factor included so that the per-neuron value is of order one under the unit-norm convention of *U*_*k*_. Averaged over the neurons of a participation-type subset *S*, it gives that subset’s normalized squared loading,

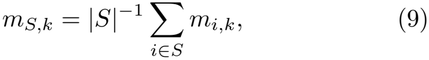

and over the entire population its mean is one by the unit-norm constraint on *U*_*k*_. The recording-wide value for each neuron is the variance-weighted average over significant dynamical PCs,

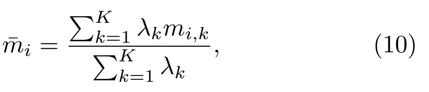

where *λ*_*k*_ is the variance of dynamical PC *k*. It follows that 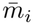 is neuron *i*’s share of the total variance of the denoised activity, relative to the average neuron.

Loading breadth distinguishes loading concentrated in a few significant dynamical PCs from loading distributed across them:

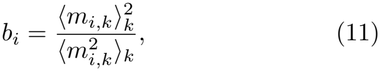

the participation ratio of a neuron’s normalized squared loadings over the *K* significant dynamical PCs. It ranges from 1*/K* when the loading is confined to one dynamical PC to one when it is equal across them. For each recording, we averaged *b*_*i*_ over the neurons in each participation type.

The dynamical variance ratio quantifies how much of the denoised collective dynamics varies on the timescale of the delay window, the burst timescale. To define it, note that the normalized total power of dynamical PC *k* decomposes as

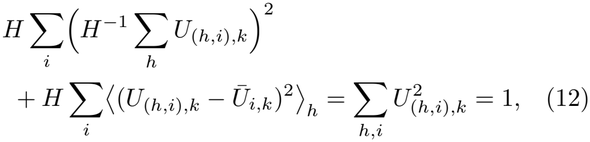

where the first term is the power 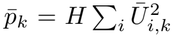 of the temporal mean profile *Ū*_*i,k*_ = *H*^−1^ ∑_*h*_ *U*_(*h,i*),*k*_ over delay channels, and the second term is the population sum 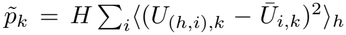 of the profile’s variance over delays. This second term, 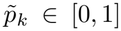, is the dynamical variance ratio of dynamical PC 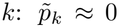 indicates a dynamical PC whose spatial profile is nearly constant over delays, reflecting slow or static collective structure, while 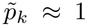 indicates a profile that varies rapidly across the delay window, reflecting dynamics on that timescale. The recording-level dynamical variance ratio is the variance-weighted mean over significant dynamical PCs,

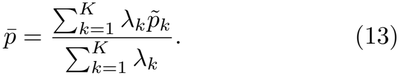

For participation-type comparisons, the dynamical variance ratio was computed over the neurons of each participation type, with dynamical PCs weighted by 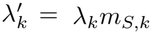, so that dynamical PCs where the type contributed little power were downweighted accordingly.

### I. Activity-Form Similarity and Clustering

Activity-form similarity compares the temporal form of each neuron’s activity within the significant dynamical PCs, independent of its normalized squared loading. For each neuron and each significant dynamical PC, the delay-resolved activity profile was divided by its rootmean-square over delays, removing the magnitude that normalized squared loading measures. The result is the neuron’s activity form in that dynamical PC. The similarity of two neurons is the variance-weighted mean, over significant dynamical PCs, of the uncentered correlation of their activity forms across delays, the cosine of the angle between the two profiles. Activity-form distance is the chord distance, 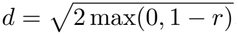, where *r* is the similarity. Chord distance was preferred over the spherical geodesic, the other common choice, because the geodesic amplifies longer distances more than shorter ones and so yields less conservative clustering.

Neurons were clustered by HDBSCAN [49, 50] applied to the activity-form distance matrix, with minimum cluster size ⌈*N/*10⌉ for a recording with *N* neurons, and neurons assigned to no cluster were treated as noise. Cluster quality was scored in the activity-form distance space by the density-based cluster-validity index (DBCV) [49], the HDBSCAN validity index, which is positive when between-cluster distances exceed within-cluster spread and negative when this ordering is reversed. The same index scored the participation-type labels on the same distance matrix, with any type of fewer than two neurons treated as noise, for the comparison in Figure 3D. For visualization, multidimensional scaling (MDS) [51] of the same distance matrix gave a two-dimensional embedding. The physical DBCV is undefined when a recording has fewer than two activity-form clusters, since no between-cluster comparison can then be made, and such recordings were excluded from that comparison.

For comparison, neurons were also clustered by their burst-to-burst latency profiles. The latency-profile distance between two neurons is the root-mean-square difference of their denoised latencies over the bursts in which both have an assigned latency, and, if they share no burst, the absolute difference of their mean latencies. HDBSCAN with the same settings was applied to this distance matrix, and the resulting DBCV was compared with the activity-form DBCV. When sufficient latency observations were available but HDBSCAN found fewer than two latency-profile clusters, this score was set to zero to represent no detected latency clustering; it was undefined and excluded only when there were insufficient usable observations.

### J. Physical-Space Analysis

Physical positions of neurons are the centroids of their CNMF spatial components. Propagation-like spatial organization within a set of neurons was quantified as the Pearson correlation between each neuron’s Euclidean distance from an early-activation reference and its latency relative to that reference. The reference position is the centroid of the two earliest-activating neurons in the set, and the reference time is their mean latency. This correlation was computed globally over all recruited neurons, those with an assigned latency in at least one burst, and separately within each non-noise activity-form cluster, where each cluster defined its own reference.

The cluster-minus-global physical correlation is the difference between a cluster’s within-cluster correlation and the global correlation of the same recording, one value per cluster. A recording with two clusters therefore contributes two values, and they are dependent. Values above zero indicate that the distance–latency organization is stronger within the cluster than in the full recruited population. Clusters with fewer than three neurons with assigned latencies were excluded.

The physical DBCV is the HDBSCAN validity index computed on the matrix of Euclidean distances between neuron positions, using the non-noise activity-form cluster labels. Values below zero indicate that neurons in the same cluster are not physically closer to one another than to neurons in other clusters.

## DATA AVAILABILITY

The calcium-imaging recordings analyzed in this study and file-placement instructions are available on Zenodo [52]. Intermediate products, including motioncorrected movies and segmentation outputs, are not deposited separately, since the supplied notebooks regenerate them from the recordings. The Zenodo archive also contains the processed plotting data needed to regenerate the figures directly, without rerunning the analyses.

## CODE AVAILABILITY

All notebooks for motion correction, segmentation, analysis, statistical testing, and figure generation are available on GitHub (https://github.com/zhaoz0314/organoid-collective-dyn, release v1.1) and archived on Zenodo [53]. Stochastic steps use fixed random seeds.

## V. APPENDIX

### A. Repeated Recordings of One Preparation Are Not Repeated Measurements

Every comparison between cytoarchitectures in the main text treats each recording as one sample, even though several preparations contribute more than one recording (Methods IV A). This appendix justifies that choice by showing that those recordings are not repeated measurements of one sample, which would otherwise call for a repeated-measures analysis of every comparison. Two considerations argue against treating them so.

First, segmentation was performed independently for each recording, and CNMF establishes no correspondence between the components extracted from different recordings of the same preparation. The same neurons were therefore not followed from one recording to the next, and two recordings of one preparation did not sample a common set of neurons, as a repeated measurement of one sample would.

Second, the data showed no preparation-level effect. Table I compares the variation between recordings of one preparation with the variation between preparations, for the three quantities where repeated recordings are available. In the CCNs, where every preparation contributed exactly one spontaneous and one excited recording, the within-preparation standard deviation exceeded the between-preparation standard deviation for all three quantities, and a one-way analysis of variance with preparation as the grouping factor detected no effect. A network’s spontaneous value also failed to predict its excited value, with correlations across the four networks of *r* = −0.07 for effective dimensionality, *r* = −0.27 for dynamical variance ratio, and *r* = −0.37 for activity-form DBCV. The BOs pointed the same way, though only two of them contributed more than one recording, so the BO evidence is weak on its own.

These null results have low power, as the BO DBCV shows. Its between-preparation standard deviation lay well above the within-preparation one, yet with only two preparations the difference fell far short of significance. The table therefore establishes only that preparationlevel structure is undetectable at this sample size. On that basis, together with the segmentation argument above, we treated each recording as one sample in every comparison in the main text.

### B. Significant Dynamical PCs Rise Well Above the Null and Account for Most of the Variance

The analyses in the main text build on the significant dynamical PCs, the components whose variance exceeds a surrogate null (Methods IV D). This appendix supports that criterion by showing the null and the spectrum it is compared with for one BO and one CCN recording, and by following the retained components of the same two recordings through their trajectories and activity forms. Across all recordings, we then report the same statistics as in the main text: the fraction of the variance those components account for, the number of dynamically distinct modes they combine into, and the share of their power that varies within single bursts.

Appendix Figure 5A and D show the dynamical PC variance spectra of the BO recording of Figure 1A and B and of an excited CCN recording, each drawn together with its surrogate null spectrum (Methods IV D). The orange lines delimit the surrogate range, the variances that finite sampling, single-neuron autocorrelation, and the delay embedding produce on their own, and its upper edge is the cutoff for significance. In both recordings, a clear gap separated the null range from the significant dynamical PCs, whose variances exceeded the edge severalfold, so their classification does not depend on the exact position of the edge.

The trajectories of the retained components (Appendix Figure 5B,E) show when each population pattern is expressed. In both example recordings, several dynamical PCs were active around the same network bursts while differing in their timescales of variation, with the lowervariance dynamical PCs varying more rapidly. The activity of individual neurons within each dynamical PC showed a similar ordering across the delay window, with faster variation in the lower-variance dynamical PCs (Appendix Figure 5C,F). Bursts therefore recruit several population patterns whose activity unfolds over different timescales.

In the example recordings, a large share of the variance lay in the significant dynamical PCs. To quantify whether this holds across recordings, we computed their summed variance divided by the summed variance of all dynamical PCs. This fraction was mostly between 50% and 80% and did not differ significantly between cytoarchitectures (BO mean 0.649, CCN mean 0.588; Welch test, *p* = 0.26; Appendix Figure 5I). Coordinated population patterns therefore account for most of the embedded variance in both cytoarchitectures.

Several significant dynamical PCs could still reflect harmonics of a single non-sinusoidal rhythm, so the dynamical PC count alone does not establish several dynamical processes. We therefore counted how many dynamically distinct modes the fitted propagator resolved (Methods IV F). Resolved frequencies were merged into one mode when all of them were distinct integer multiples, up to the sixth, of a common fundamental within 5%, and each real eigenvalue, which labels a nonoscillating mode, counted as one further mode. In the BO example, the three resolved frequencies were not integer multiples of one another, and in the CCN example a single oscillation coexisted with one non-oscillating mode (Appendix Figure 5G,H). Across recordings, the number of distinct modes exceeded one in both cytoarchitectures (BO: mean 2.11, *p*_Holm_ = 0.013; CCN: mean 1.62, *p*_Holm_ = 0.025), and the two were not distinguishable from each other (Welch test, *p* = 0.29; Appendix Figure 5K). Effective dimensionality therefore separates the cytoarchitectures where the number of distinct dynamical modes does not, so the larger BO repertoire consists of more dynamical components distributed over a similar set of oscillations.

The main text uses the dynamical variance ratio to ask whether denoised activity changes within single bursts, and here we report its value for whole recordings. The ratio (Methods IV H) is the share of a dynamical PC’s power that varies across the delay window, near zero for a component whose population pattern stays nearly constant over a burst and near one for a component whose pattern changes within it. Averaged over the significant dynamical PCs of each recording, this ratio was significantly greater than zero in both cytoarchitectures (BO: mean 0.336, *p*_Holm_ = 1.2× 10^−4^; CCN: mean 0.211, *p*_Holm_ = 0.0030) and did not differ significantly between them (Welch test, *p* = 0.11; Appendix Figure 5J). Both cytoarchitectures therefore contain population patterns that change within single bursts.

### C. Pharmacological Excitation Increases Bursting Without Altering Other Measured Properties of Denoised Dynamics

The main text pools spontaneous and pharmacologically excited recordings in every comparison between cytoarchitectures, on the grounds that excitation changes how much activity a network produces while leaving its organization unchanged. This appendix documents both halves of that claim, starting from the known effect of excitation on the rate of network activity [22]. Appendix Figure 6A shows the CNMF segmentation of an example CCN under the two conditions, and Appendix Figure 6B its activity, with bursts visibly more frequent under excitation. Across all spontaneous and excited recordings, including those with no detected bursts, the burst rate rose from a spontaneous mean of 0.68 to an excited mean of 2.55 bursts per minute (Appendix Figure 6C; *n* = 14 and 11 recordings, respectively; one-sided Welch t-test, *p* = 0.035). Muscimol recordings were excluded from both groups.

The remaining measures require multiple bursts, so the two conditions were compared among the 17 multi-burst recordings, with a two-way analysis of variance for condition, cytoarchitecture, and their interaction, for the measures of denoised dynamics listed in Table II. No condition term was significant. Appendix Figure 6D–G show four of these measures: effective dimensionality (Figure 1F), dynamical variance ratio (Appendix Figure 5J), order stability (Appendix Figure 7B), and activity-form DBCV (Figure 3C). Excitation therefore makes population bursts more frequent while leaving their organization, as far as we measured it, unchanged, and this is what allows the two conditions to be pooled.

**TABLE II.** Two-Way ANOVA Across Measures of Denoised Dynamics. Uncorrected *p* values for the condition term (spontaneous vs. pharmacologically excited), the cytoar-chitecture term (BO vs. CCN), and their interaction among the 17 multi-burst recordings; all rows use one value per recording except within-type gain, which uses one value per activity type. Bold marks *p* < 0.05. No condition term is significant. The cytoarchitecture term for effective dimensionality corresponds to the difference reported in the main text; the other significant cytoarchitecture terms do not survive Bon-ferroni correction across the ten cytoarchitecture tests and are not discussed elsewhere in the paper.

| quantity | condition | cytoarch. | interaction |
|---|---|---|---|
| effective dimensionality | 0.70 | **0.006** | 0.55 |
| significant dynamical PC variance fraction | 0.22 | 0.27 | 0.13 |
| dynamical variance ratio | 0.28 | 0.13 | 0.78 |
| fundamental frequency | 0.52 | 0.64 | 0.75 |
| order stability | 0.11 | **0.019** | 0.76 |
| activity-form DBCV | 0.16 | 0.053 | 0.15 |
| number of activity types | 0.26 | 0.083 | 0.098 |
| physical DBCV | 0.058 | 0.42 | **0.039** |
| within-type gain in distance–latency correlation | 0.069 | 0.27 | 0.30 |
| global distance–latency correlation | 0.059 | **0.010** | 0.87 |

One interaction term in Table II was significant, for the physical DBCV. Ten interaction terms were tested without correction, so one value below 0.05 is expected by chance alone, and the interaction concerned differences among scores that all lay well below zero, where the activity types are not segregated in space in either cytoarchitecture or condition. It therefore does not affect the pooling of conditions.

### D. Denoising Preserves Activation Order

The main text introduces the activation sequence on raw CNMF activity (Figure 2A) and then measures latencies, participation, and order on denoised activity (Figure 2B onward). This appendix checks across recordings that denoising preserves the mean activation order while making it more stable from burst to burst.

Before denoising, there is no residual fluctuation against which to judge whether a neuron activates, so the raw readout of Figure 2A assigned a latency in every burst window that contained a peak and was otherwise computed in the same way (Methods IV G). In the latency matrix of Figure 2A-right, the colors along each row varied more from burst to burst than in Figure 2B-right, but neurons that were early on average tended to stay early, so the order was already visible before denoising.

To check that denoising keeps this order, we measured the Kendall similarity between the mean CNMF latencies and the mean denoised latencies of the neurons in each recording (Appendix Figure 7A). It was high across both cytoarchitectures, with a median of 0.65 and 13 of 17 recordings above 0.4, and it did not differ significantly between them (Welch test, BO vs. CCN: *p* = 0.84). The remaining four recordings lay near or below zero, two of them negative and not shown in the panel, so in those the mean order was not preserved through denoising. Denoising therefore preserves the mean activation order in most recordings.

Denoising should also make the order more stable from burst to burst, since the significant dynamical PCs keep the parts of the activity that repeat across bursts. Within each recording, we averaged the Kendall similarity of the latencies over all pairs of bursts, using for each pair every neuron with an assigned latency in both bursts (Methods IV G), so that higher values mean the relative order repeats more closely from burst to burst. Appendix Figure 7B compares this order stability before and after denoising within each recording, with gray lines connecting the same recording. Denoised activity had greater order stability than CNMF activity (paired test, *p* = 2.2 × 10^−7^), as expected.

### E. Significant Dynamical PC Loading Extends Beyond Sequence Participants

The main text reports that neurons of all three participation types load on the denoised activity, and that significant dynamical PCs occur even in recordings that never burst. Loading above zero is a weak criterion, so here we measure how broadly each participation type spreads its loading over the significant dynamical PCs, test whether the dynamical PCs would remain significant if only one participation type contributed, and compare the significant dynamical PCs of non-bursting recordings with those of the non-participating neurons in bursting ones.

Appendix Figure 8A plots the normalized squared loading of every neuron in each significant dynamical PC of the CCN recording of Figure 2, sorted within each dynamical PC and colored by participation type. Neurons of all three types appeared throughout the range of loading, and the mean loading of each type (dashed lines) changed from one dynamical PC to the next.

A neuron’s contribution to the denoised dynamics can be concentrated in one significant dynamical PC or distributed across several. We distinguished these possibilities with the loading breadth (Methods IV H), the participation ratio of each neuron’s normalized squared loading across dynamical PCs, averaged within each participation type for every recording (Appendix Figure 8B). Consistent neurons distributed their loading more broadly across dynamical PCs than intermittent and non-participating neurons, while the latter two types did not differ (means across recordings: consistent 0.72, intermittent 0.50, non-participating 0.54; Friedman test, *p* = 1.8 ×10^−4^; Holm-corrected paired tests, consistent vs. intermittent and non-participating, both *p*_Holm_ = 5.5×10^−4^; intermittent vs. non-participating, *p*_Holm_ = 0.38; *n* = 15 recordings containing all three types). Neurons that join every burst therefore contribute across more of the collective repertoire, while neurons that join intermittently or never concentrate their loading in fewer dynamical PCs.

To test whether one participation type alone would support the collective patterns, we adjusted each dynamical PC’s variance by the mean normalized squared loading of that type in that dynamical PC (Methods IV H), which estimates the variance the dynamical PC would have if every neuron loaded on it like a neuron of that type. Comparing the adjusted variance with the upper edge of the surrogate spectrum then tells whether the dynamical PC would still be significant. In the same example recording, two of the three significant dynamical PCs stayed above the edge for the non-participating and the intermittent neurons, and all three for the consistent neurons (Appendix Figure 8C). Across recordings, the adjusted-significance fraction, the fraction of a recording’s significant dynamical PCs that stay above the edge after adjustment, was greater than zero for all three types (one-sample tests, all *p*_Holm_ < 10^−6^; means: non-participating 0.45, intermittent 0.53, consistent 0.93), higher for consistent neurons than for the other two, and indistinguishable between intermittent and non-participating neurons (ANOVA, *p* = 1.1 ×10^−7^; consistent vs. intermittent, *p*_Holm_ = 3.6× 10^−5^; consistent vs. non-participating, *p*_Holm_ = 2.4 ×10^−7^; intermittent vs. non-participating, *p*_Holm_ = 0.32; Appendix Figure 8D). About half of the significant dynamical PCs would therefore remain significant on the loading of the non-participating neurons alone, which is the sense in which these neurons remain part of the denoised activity.

Ten recordings showed no bursts at all (Methods IV A), and their collective activity, if any, should resemble what the non-participating neurons express in bursting recordings. Non-bursting recordings did have significant dynamical PCs, on average more of them than remained significant in bursting recordings after adjustment to the loading of the non-participating neurons (means 2.6 and 1.8; Welch test, *p* = 0.027; Appendix Figure 8E). In both, the dynamical variance ratio was close to zero, not significantly above it for the non-bursting recordings and only weakly for the non-participating neurons, and the two did not differ (one-sample tests against zero: non-bursting *p*_Holm_ = 0.056, non-participating *p*_Holm_ = 0.021; Welch test, *p*_Holm_ = 0.10; Appendix Figure 8F). Non-bursting recordings therefore contain significant collective components, and these vary on timescales slower than a burst, like the activity of the non-participating neurons in bursting recordings.

### F. Consistent Neurons Vary Widely in the Amount of Short-Timescale Dynamics

The main text reports that the dynamical variance ratio of consistent neurons is higher on average than that of the other two participation types but varies widely across recordings, approaching the other types in some, and it calls the values a single continuum. The alternative is two groups of recordings, in one of which the consistent neurons express burst-timescale dynamics and in the other not. Appendix Figure 9 shows the distribution of the consistent-neuron ratio across the multiburst recordings, with each recording marked by condition and cytoarchitecture. A Gaussian-mixture comparison favored one component over two (ΔBIC_1−2_ = −7.5), so the spread is one broad distribution and does not resolve into two groups of recordings.

### G. The Number of Activity Types Is Similar Across Cytoarchitectures

The main text reports that activity types are similarly separated in BOs and CCNs (Figure 3C), with typically two types per recording. The number of types could still differ between cytoarchitectures. Appendix Figure 10A and B show the distribution of the number of activity types per recording in each cytoarchitecture, and Appendix Figure 10C compares them: the median was two in both, and the difference was not significant (Welch test, *p* = 0.087). The similar separation in the main text therefore does not hide a difference in how many activity types each cytoarchitecture supports.

### H. Activity Types Are Separated by High-Variance, Slow Significant Dynamical PCs

The main text reports that the activity types are separated mostly along one strong, slow component of the collective activity. In the BO recording of Figure 3, the centroid activity forms of the two types took opposite signs on one nearly flat dynamical PC and overlapped on the others (Appendix Figure 11A). To test whether this holds across recordings, we took the principal axes of the spread of the type centroids in standardized activity-form space, one fewer than the number of types, and measured the share of each axis that fell on each significant dynamical PC. Pooled over recordings, axes, and dynamical PCs, this share increased with the variance of the dynamical PC (Pearson *r* = 0.55, *p* = 9.7× 10^−8^; Appendix Figure 11B) and decreased with its dynamical variance ratio (Pearson *r* = −0.48, *p* = 3.9× 10^−6^; Appendix Figure 11C). The activity types are therefore separated by strong, slowly varying dynamical PCs, which is why they need not follow sequence participation, a division on the faster burst timescale.

## ACKNOWLEDGEMENTS

We thank Gordon Berman, Tankut Can, Srdjan Ostojic, Flora Vaccarino, and Soraya Scuderi for thoughtful feedback. I.N. and Z.Z. were supported in part by a Simons Foundation Investigator grant and by NIH grants R01-NS084844 and R01-NS099375. L.Y. and A.L. were supported by NIH grant RF1-MH123978 and a Yale Kavli Institute for Neuroscience Innovator Award.

## AUTHOR CONTRIBUTIONS

Z.Z. performed the analysis. L.Y. and A.L. led the experimental work. I.N. led the analysis. Z.Z., A.L., and I.N. wrote the paper. A.L. and I.N. secured funding and are co-corresponding authors.

## References

[1] J. B. Ackman, T. J. Burbridge, and M. C. Crair, Retinal waves coordinate patterned activity throughout the developing visual system, Nature 490, 219 (2012).

[2] M. W. Wu, N. Kourdougli, and C. Portera-Cailliau, Network state transitions during cortical development, Nat. Rev. Neurosci. 25, 535 (2024).

[3] J. P. Cunningham and B. M. Yu, Dimensionality reduction for large-scale neural recordings, Nat. Neurosci. 17, 1500 (2014).

[4] K. V. Shenoy, M. Sahani, and M. M. Churchland, Cortical control of arm movements: a dynamical systems perspective, Annu. Rev. Neurosci. 36, 337 (2013).

[5] E. Pastalkova, V. Itskov, A. Amarasingham, and G. Buzsáki, Internally generated cell assembly sequences in the rat hippocampus, Science 321, 1322 (2008).

[6] A. P. Vaz, J. H. Wittig, S. K. Inati, and K. A. Zaghloul, Backbone spiking sequence as a basis for preplay, replay, and default states in human cortex, Nat. Commun. 14, 4723 (2023).

[7] M. A. Lancaster, M. Renner, C.-A. Martin, D. Wenzel, L. S. Bicknell, M. E. Hurles, T. Homfray, J. M. Penninger, A. P. Jackson, and J. A. Knoblich, Cerebral organoids model human brain development and microcephaly, Nature 501, 373 (2013).

[8] T. Kadoshima, H. Sakaguchi, T. Nakano, M. Soen, S. Ando, M. Eiraku, and Y. Sasai, Self-organization of axial polarity, inside-out layer pattern, and species-specific progenitor dynamics in human ES cell-derived neocortex, Proc. Natl. Acad. Sci. U.S.A. 110, 20284 (2013).

[9] M. A. Lancaster, N. S. Corsini, S. Wolfinger, E. H. Gustafson, A. W. Phillips, T. R. Burkard, T. Otani, F. J. Livesey, and J. A. Knoblich, Guided self-organization and cortical plate formation in human brain organoids, Nat. Biotechnol. 35, 659 (2017).

[10] X. Qian, H. Song, and G.-l. Ming, Brain organoids: Advances, applications and challenges, Development 146, dev166074 (2019).

[11] G. Quadrato, T. Nguyen, E. Z. Macosko, J. L. Sherwood, S. M. Yang, D. R. Berger, N. Maria, J. Scholvin, M. Gold-man, J. P. Kinney, E. S. Boyden, J. W. Lichtman, Z. M. Williams, S. A. McCarroll, and P. Arlotta, Cell diversity and network dynamics in photosensitive human brain organoids, Nature 545, 48 (2017).

[12] Z. Zhang, R. O’Laughlin, H. Song, and G.-l. Ming, Patterning of brain organoids derived from human pluripotent stem cells, Current Opinion in Neurobiology 74, 102536 (2022).

[13] A. Amiri, G. Coppola, S. Scuderi, F. Wu, T. Roychowdhury, F. Liu, S. Pochareddy, Y. Shin, A. Safi, L. Song, Y. Zhu, A. M. M. Sousa, PsychENCODE Consortium, M. Gerstein, G. E. Crawford, N. Sestan, A. Abyzov, and F. M. Vaccarino, Transcriptome and epigenome landscape of human cortical development modeled in organoids, Science 362, eaat6720 (2018).

[14] S. Scuderi, T. Y. Kang, A. Jourdon, A. Nelson, L. Yang, F. Wu, G. M. Anderson, J. Mariani, L. Tomasini, V. Sarangi, A. Abyzov, A. Levchenko, and F. M. Vaccarino, Specification of human brain regions with orthogonal gradients of WNT and SHH in organoids reveals patterning variations across cell lines, Cell Stem Cell 32, 970 (2025).

[15] J. Mariani, M. V. Simonini, D. Palejev, L. Tomasini, G. Coppola, A. M. Szekely, T. L. Horvath, and F. M. Vaccarino, Modeling human cortical development in vitro using induced pluripotent stem cells, Proc. Natl. Acad. Sci. U.S.A. 109, 12770 (2012).

[16] J. Mariani, G. Coppola, P. Zhang, A. Abyzov, L. Provini, L. Tomasini, M. Amenduni, A. Szekely, D. Palejev, M. Wilson, M. Gerstein, E. L. Grigorenko, K. Chawarska, K. A. Pelphrey, J. R. Howe, and F. M. Vaccarino, FOXG1-dependent dysregulation of GABA/glutamate neuron differentiation in autism spectrum disorders, Cell 162, 375 (2015).

[17] E. Mire, C. Mezzera, E. Leyva-Díaz, A. V. Paternain, P. Squarzoni, L. Bluy, M. Castillo-Paterna, M. J. López, S. Peregrín, M. Tessier-Lavigne, S. Garel, J. Galcerán, J. Lerma, and G. López-Bendito, Spontaneous activity regulates Robo1 transcription to mediate a switch in thalamocortical axon growth, Nat. Neurosci. 15, 1134 (2012).

[18] S. A. Sloan, J. Andersen, A. M. Paşca, F. Birey, and S. P. Paşca, Generation and assembly of human brain region-specific three-dimensional cultures, Nat. Protoc. 13, 2062 (2018).

[19] H. Sakaguchi, Y. Ozaki, T. Ashida, T. Matsubara, N. Oishi, S. Kihara, and J. Takahashi, Self-organized synchronous calcium transients in a cultured human neural network derived from cerebral organoids, Stem Cell Rep. 13, 458 (2019).

[20] K. J. Brennand, A. Simone, J. Jou, C. Gelboin-Burkhart, N. Tran, S. Sangar, Y. Li, Y. Mu, G. Chen, D. Yu, S. Mc-Carthy, J. Sebat, and F. H. Gage, Modelling schizophrenia using human induced pluripotent stem cells, Nature 473, 221 (2011).

[21] C. Stringer, M. Pachitariu, N. Steinmetz, C. B. Reddy, M. Carandini, and K. D. Harris, Spontaneous behaviors drive multidimensional, brainwide activity, Science 364, eaav7893 (2019).

[22] C. A. Trujillo, R. Gao, P. D. Negraes, J. Gu, J. Buchanan, S. Preissl, A. Wang, W. Wu, G. G. Haddad, I. A. Chaim, A. Domissy, M. Vandenberghe, A. Devor, G. W. Yeo, B. Voytek, and A. R. Muotri, Complex oscillatory waves emerging from cortical organoids model early human brain network development, Cell Stem Cell 25, 558 (2019).

[23] T. van der Molen, A. Spaeth, M. Chini, S. Hernandez, G. A. Kaurala, H. E. Schweiger, C. Duncan, S. McKenna, J. Geng, M. Lim, J. Bartram, T. Gänswein, A. Dendukuri, Z. Zhang, J. Gonzalez-Ferrer, K. Bhaskaran-Nair, A. L. Morson, C. R. K. Harder, L. R. Petzold, D.-M. Alam El Din, J. Laird, M. Schenke, L. Smirnova, B. M. Colquitt, M. A. Mostajo-Radji, P. K. Hansma, M. Teodorescu, A. Hierlemann, K. B. Hengen, I. L. Hanganu-Opatz, K. S. Kosik, and T. Sharf, Preconfigured neuronal firing sequences in human brain organoids, Nat. Neurosci. 29, 123 (2026).

[24] E. A. Pnevmatikakis, D. Soudry, Y. Gao, T. A. Machado, J. Merel, D. Pfau, T. Reardon, Y. Mu, C. Lacefield, W. Yang, M. Ahrens, R. Bruno, T. M. Jessell, D. S. Peterka, R. Yuste, and L. Paninski, Simultaneous denoising, deconvolution, and demixing of calcium imaging data, Neuron 89, 285 (2016).

[25] A. Giovannucci, J. Friedrich, P. Gunn, J. Kalfon, B. L. Brown, S. A. Koay, J. Taxidis, F. Najafi, J. L. Gauthier, P. Zhou, B. S. Khakh, D. W. Tank, D. B. Chklovskii, and E. A. Pnevmatikakis, CaImAn an open source tool for scalable calcium imaging data analysis, eLife 8, e38173 (2019).

[26] S. L. Brunton, B. W. Brunton, J. L. Proctor, E. Kaiser, and J. N. Kutz, Chaos as an intermittently forced linear system, Nat. Commun. 8, 19 (2017).

[27] M. Potters and J.-P. Bouchaud, A First Course in Random Matrix Theory: for Physicists, Engineers and Data Scientists (Cambridge University Press, Cambridge, 2020).

[28] R. J. Bell and P. Dean, Atomic vibrations in vitreous silica, Discuss. Faraday Soc. 50, 55 (1970).

[29] H. Ko, S. B. Hofer, B. Pichler, K. A. Buchanan, P. J. Sjöström, and T. D. Mrsic-Flogel, Functional specificity of local synaptic connections in neocortical networks, Nature 473, 87 (2011).

[30] C. Holmgren, T. Harkany, B. Svennenfors, and Y. Zilberter, Pyramidal cell communication within local networks in layer 2/3 of rat neocortex, The Journal of Physiology 551, 139 (2003).

[31] R. Perin, T. K. Berger, and H. Markram, A synaptic organizing principle for cortical neuronal groups, Proc. Natl. Acad. Sci. U.S.A. 108, 5419 (2011).

[32] J. Peng, A. Duffy, I. Dippenaar, L. Liu, J. Wu, X. Tu, A. L. Fairhall, and C. Lois, Flexible neuronal participation within a reliable motor sequence, bioRxiv 10.64898/2026.07.10.737866 (2026).

[33] M. Diesmann, M.-O. Gewaltig, and A. Aertsen, Stable propagation of synchronous spiking in cortical neural networks, Nature 402, 529 (1999).

[34] R. Cheong, A. Rhee, C. J. Wang, I. Nemenman, and A. Levchenko, Information transduction capacity of noisy biochemical signaling networks, Science 334, 354 (2011).

[35] A. Mugler, A. Levchenko, and I. Nemenman, Limits to the precision of gradient sensing with spatial communication and temporal integration, Proc. Natl. Acad. Sci. U.S.A. 113, E689 (2016).

[36] D. Ellison, A. Mugler, M. D. Brennan, S. H. Lee, R. J. Huebner, E. R. Shamir, L. A. Woo, J. Kim, P. Amar, I. Nemenman, A. J. Ewald, and A. Levchenko, Cell–cell communication enhances the capacity of cell ensembles to sense shallow gradients during morphogenesis, Proc. Natl. Acad. Sci. U.S.A. 113, E679 (2016).

[37] I. Nemenman and P. Mehta, Random-with-constraints: Constructing minimal models for high-dimensional biology, Proc. Natl. Acad. Sci. U.S.A. 123, e2526777123 (2026).

[38] D. G. Clark, L. F. Abbott, and A. Litwin-Kumar, Dimension of activity in random neural networks, Phys. Rev. Lett. 131, 118401 (2023).

[39] Z. Zhao, M. J. Pasek, and I. M. Nemenman, Random neural networks match observed dimensionality of neural population recordings and motivate stronger experimental tests, arXiv:2605.26551 [q-bio.NC] (2026), arXiv:2605.26551 [q-bio.NC].

[40] J. Kim, J. A. Efe, S. Zhu, M. Talantova, X. Yuan, S. Wang, S. A. Lipton, K. Zhang, and S. Ding, Direct reprogramming of mouse fibroblasts to neural progenitors, Proc. Natl. Acad. Sci. U.S.A. 108, 7838 (2011).

[41] Z. Burda, J. Jurkiewicz, and B. Waclaw, Spectral moments of correlated Wishart matrices, Phys. Rev. E 71, 026111 (2005).

[42] G. E. P. Box, G. M. Jenkins, and G. C. Reinsel, Time Series Analysis: Forecasting and Control, 4th ed. (Wiley, 2008).

[43] L. Mazzucato, A. Fontanini, and G. La Camera, Stimuli reduce the dimensionality of cortical activity, Front. Syst. Neurosci. 10, 11 (2016).

[44] P. Gao, E. Trautmann, B. Yu, G. Santhanam, S. Ryu, K. Shenoy, and S. Ganguli, A theory of multineuronal dimensionality, dynamics and measurement, bioRxiv:214262 [q-bio.NC] (2017), bioRxiv:214262.

[45] P. J. Schmid, Dynamic mode decomposition of numerical and experimental data, J. Fluid Mech. 656, 5 (2010).

[46] J. N. Kutz, S. L. Brunton, B. W. Brunton, and J. L. Proctor, Dynamic Mode Decomposition: Data-Driven Modeling of Complex Systems (Society for Industrial and Applied Mathematics, Philadelphia, 2016).

[47] S. T. M. Dawson, M. S. Hemati, M. O. Williams, and C. W. Rowley, Characterizing and correcting for the effect of sensor noise in the dynamic mode decomposition, Exp. Fluids 57, 42 (2016).

[48] M. G. Kendall, A new measure of rank correlation, Biometrika 30, 81 (1938).

[49] R. J. G. B. Campello, D. Moulavi, A. Zimek, and J. Sander, Hierarchical density estimates for data clustering, visualization, and outlier detection, ACM Trans. Knowl. Discov. Data 10, 5:1 (2015).

[50] L. McInnes, J. Healy, and S. Astels, hdbscan: Hierarchical density based clustering, J. Open Source Softw. 2, 205 (2017).

[51] J. B. Kruskal, Multidimensional scaling by optimizing goodness of fit to a nonmetric hypothesis, Psychometrika 29, 1 (1964).

[52] Z. Zhao, T.-Y. Kang, L. Yang, I. Nemenman, and A. Levchenko, Calcium-imaging recordings from brain organoids (BOs) and cable-cluster networks (CCNs) (2026).

[53] Z. Zhao, zhaoz0314/organoid-collective-dyn: Analysis code for collective dynamics in BOs and CCNs, Zenodo, version v1.1, doi:10.5281/zenodo.24102516 (2026).

